# Epithelial Self-Organization Patterns the Tadpole Oral Disc

**DOI:** 10.64898/2026.10.11.758282

**Authors:** Cheng Chen, Nuo Ding, Ying Sun, Jiaxing Huang, Shuo Wang

## Abstract

Vertebrate epidermal organs are generally patterned through epithelial-mesenchymal interactions, but whether such interactions are essential for all epidermal organs remains unclear. Here we show that the elaborate oral disc of anuran tadpoles arises by partitioning a continuous oral epithelium into developmentally distinct jaw, tooth-row and papilla modules, in the absence of a mesenchymal organizer. Spatially restricted epithelial programs specify, maintain and differentiate these modules, enabling their semi-independent development. Perturbation of *lef1*, *hoxc13*, *sox7* and *pitx2* selectively disrupts individual modules. The resulting module-specific reductions, expansions and rearrangements mirror oral-disc configurations that have evolves repeatedly across anurans. These findings indicate that differential deployment of shared epithelial programs can generates selectable variation in the tadpole oral disc, allowing transient larval feeding structures to diversify independently of adult ecology and enabling tadpoles to exploit diverse ecological niches. Epidermal self-organization can therefore generate structural complexity, promote functional diversification and enhance evolvability in transient vertebrate organs.

## Introduction

How complex and spatially organized structures arise from initially continuous tissues is a fundamental problem in animal development. In vertebrates, epidermal appendages are generally considered to be induced, patterned and maintained by reciprocal signaling between the epithelium and the underlying mesenchyme(*1*). Such epithelial–mesenchymal interactions direct the development of diverse integumentary derivatives including hair follicles(*1*), feathers(*2*), teeth(*3*) and cutaneous glands(*4*), and provide the conceptual basis for prevailing models of epidermal-organ homeostasis and regeneration(*5*). Whether complex epithelial patterns can arise when this canonical tissue interaction is absent remains much less clear.

Anuran tadpoles present a challenge to this canonical framework. Across the tadpole body, the epidermis rests directly on an acellular collagenous lamella without an intervening cellular dermis, even at developmental stages when specialized epidermal structures first emerge(*6*). The formation of these structures within this tissue context raises the possibility that some vertebrate epithelia possess a substantially greater intrinsic capacity for induction and pattern formation than canonical epithelial-mesenchymal models predict.

The tadpole oral disc offers a tractable system for testing this possibility. This elaborate, transient larval feeding organ comprises several morphologically and functionally distinct epithelial structures, including keratinized jaws, labial tooth rows and marginal papillae(*7*). These structures occupy stereotyped positions within the oral field, yet their number, morphology and arrangement vary substantially among anuran species, often in association with variation in habitat, diet and feeding mode(*7, 8*). Although oral-disc characters have long been used in tadpole taxonomy, ecology and evolutionary studies(*7, 8*), the mechanism by which a continuous oral epithelium is partitioned into discrete, species-specific structural domains remains poorly understood, particularly given the apparent absence of a canonical dermal organizer(*9*).

Here we compared oral development in the Asian toad *Bufo gargarizans*, which forms a typical oral disc, with that in the African clawed frog *Xenopus laevis*, which lacks this characteristic larval structure. We traced the divergence of their oral epithelia after mouth opening and mapped how the continuous oral epithelium of *B. gargarizans* is partitioned into distinct territories that generate the jaws, tooth rows and marginal papillae. We then defined the tissue context in which this patterning occurs, identified candidate regulators and tested their roles in specifying individual epithelial modules. We further asked whether these modules arise without a morphologically discernible cellular mesenchymal organizer, thereby probing the capacity of the oral epithelium to generate complex spatial patterns intrinsically. Our findings establish the tadpole oral disc as an experimentally tractable model of epithelium-intrinsic patterning and identify a developmental route by which complex vertebrate epidermal structures can arise in the absence of a morphologically distinct mesenchymal organizer.

## Results

### Oral epithelial morphogenesis diverges after mouth opening

*B. gargarizans* tadpoles developed a typical oral disc encircling a centrally positioned mouth, with keratinized upper and lower jaw sheaths, two anterior and three posterior labial tooth rows, and non-keratinized marginal and submarginal papillae (Fig. 1A-D). By contrast, *X. laevis* lacked these perioral structures (Fig. 1E-H). Before primary mouth opening, the oral tissues of both species expanded outwards and contained BrdU-positive basal cells together with widespread E-cadherin expression, consistent with a predominantly epithelial organization (Figs. 1I, S1-3). Following mouth opening, the oral epithelium continued to expand as the maxillary and mandibular cartilages developed in both species (Fig. 1I-J). Subsequent epithelial differentiation, however, followed sharply divergent trajectories.

**Fig. 1.**
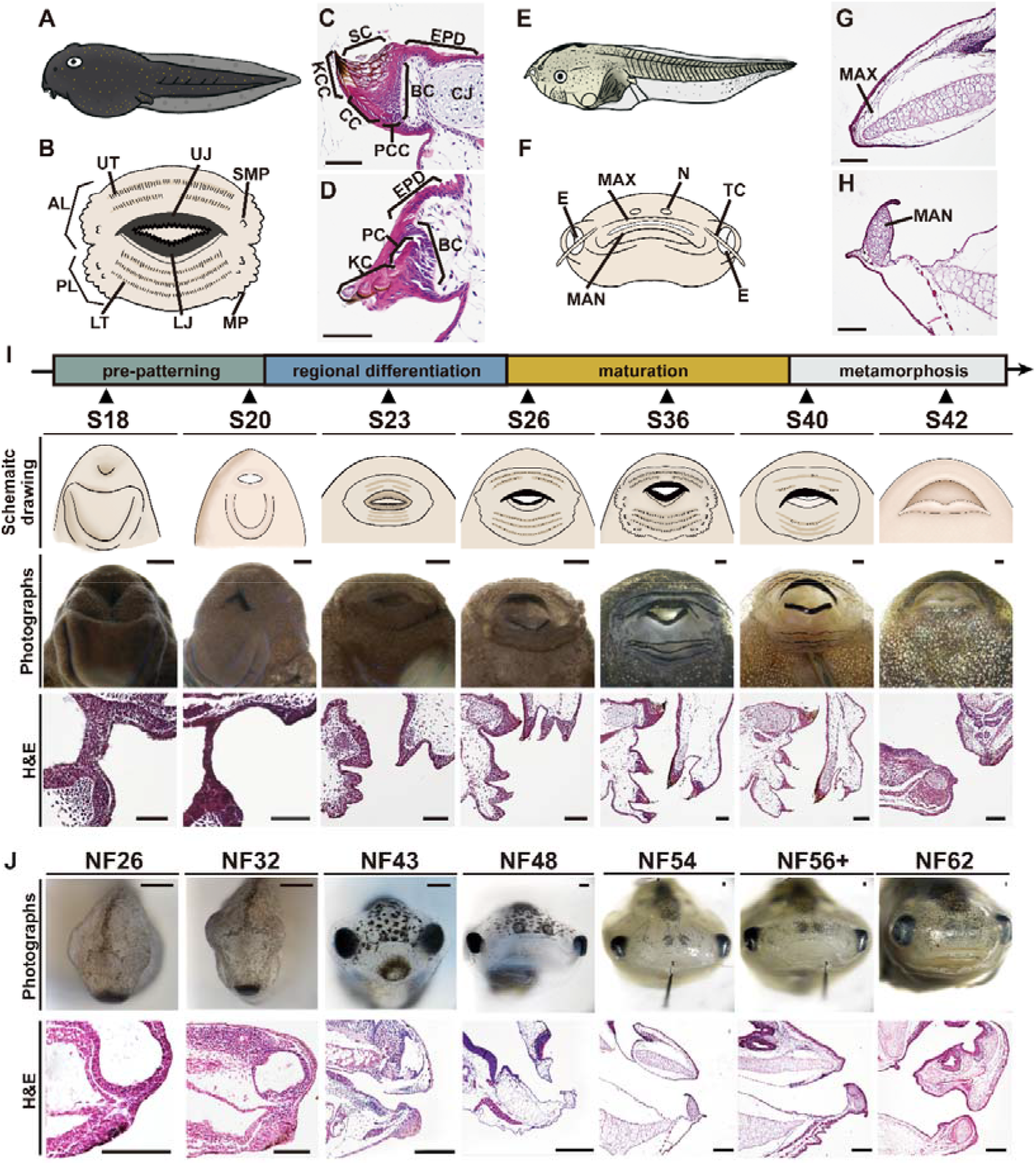
Structural organization and developmental dynamics of tadpole oral structures in *B. gargarizans* and *X. laevis*. A, Schematic illustrations of *B. gargarizans* tadpoles in left lateral view, showing the anteroventral orientation of the mouth. **Not to scale.** **B,** Schematic illustration of the oral disc of *B. gargarizans*. Jaw sheaths and labial tooth rows constitute the principal keratinized feeding structures, whereas marginal and submarginal papillae are non-keratinized epithelial projections. **Not to scale**. **C-D,** H&E staining of the jaw (**C**) and labial tooth row (**D**) of *B. gargarizans* tadpoles at S36 (sagittal sections). The jaw forms a compact triangular epithelial structure with an inwardly curved keratinized edge supported by loose connective tissue attached to the jaw cartilage (CJ). Basal precone cells on the labial side flatten into cone cells during keratinization, forming overlapping shingle-like layers that generate the serrated cutting edge, whereas lingual sheath cells retain nuclei and enlarge. Labial tooth rows are slender epithelial ridges in which teeth differentiate from proliferative basal cells and progressively keratinize into lingually curved cusps. No detectable dermis is present beneath the specialized oral epithelium. **Scale bars= 75**μ**m.** **E,** Schematic illustration of a *Xenopus laevis* tadpole in left lateral view, showing the anteroventral orientation of the mouth. **Not to scale.** **F,** Schematic illustration of the oral region of *X. laevis*, showing the absence of a typical oral disc. Protrusions adjacent to the tentacles may represent papilla-like structures, although their homology to oral-disc papillae remains unresolved. **Not to scale.** **G-H,** H&E staining of the upper (**G**) and lower (**H**) jaws of *X. laevis* tadpoles at NF54 (sagittal sections). The mouth is supported directly by the maxillary and mandibular cartilages and lacks oral-disc-associated keratinized feeding structures. No dermis is detected beneath the oral epithelium. **Scale bars= 150**μ**m.** **I,** Morphological and histological development of the *B. gargarizans* oral disc across seven representative stages spanning four phases: pre-patterning, regional differentiation, maturation, and metamorphic regression. Panels show external morphology and corresponding H&E staining. Jaw sheaths and labial tooth rows become morphologically distinct during regional differentiation, keratinize during maturation, and regress during metamorphosis. **Scale bars= 250**μ**m.** **J.** Development of the oral region in *X. laevis* from NF26 to NF62 corresponding to the representative stages in *Bufo*, showing gross morphology and histological features of the oral apparatus. The presumptive mouth and oral depression become apparent early in development, whereas keratinized oral-disc structures are absent throughout. As development proceeds, the oral margin expands and the maxillary and mandibular cartilages differentiate, but jaw sheaths, labial tooth rows, and marginal papillae do not form. **Scale bars= 250**μ**m.** **Abbreviations: AL**, anterior labium; **BC,** basal cell; **CC,** cone cell; **CJ,** jaw cartilage; **E,** eye; **EPD,** non-keratinized epidermis; **KC,** keratinized cusp; **KCC,** keratinized cone cell; **LJ,** lower jaw sheath; **LT,** lower tooth row; **MAN,** mandibular cartilage; **MAX,** maxillary cartilage; **MP,** marginal papillae; **N,** nostril; **PC,** precusp cell; **PCC,** precone cell; **PL,** posterior labium; **SC,** sheath cell; **SMP,** submarginal papillae; **TC,** tentacle; **UJ,** upper jaw sheath; **UT,** upper tooth row.

In *B. gargarizans*, the oral epithelium progressively thickened and folded within the prospective jaw and tooth-row territories (Fig. 1I, Fig. S1A). At S23, HCR detected *vimentin* transcripts in elongating cells at the leading edge of the jaw epithelium, with little expression in the tooth-row epithelium (Fig. S2C), whereas vimentin protein remained undetectable by immunofluorescence (Fig. 2A; Fig. S2C). By S26, vimentin protein was detectable in both jaw and tooth-row epithelia, accompanying sheath-cell elongation and precusp-cell differentiation (Fig. 2A; Fig. S2C). Distal keratinization also became evident by S26, and continued epithelial growth and regional differentiation produced a mature oral disc by S36 (Fig. 1I). In the mature oral disc, the jaw sheaths consisted of asymmetric stratified epithelia, in which cone and sheath cells formed an inwardly curved keratinized cutting edge (Fig. 1C). The tooth ridges generated lingually curved keratinized cusps (Fig. 1D), whereas the marginal papillae persisted as non-keratinized epithelial projections (Fig. 1I). Thus, a continuous oral epithelium gave rise to distinct jaw, tooth-row and papillary architectures, defining these territories as discrete morphogenetic modules.

**Fig. 2.**
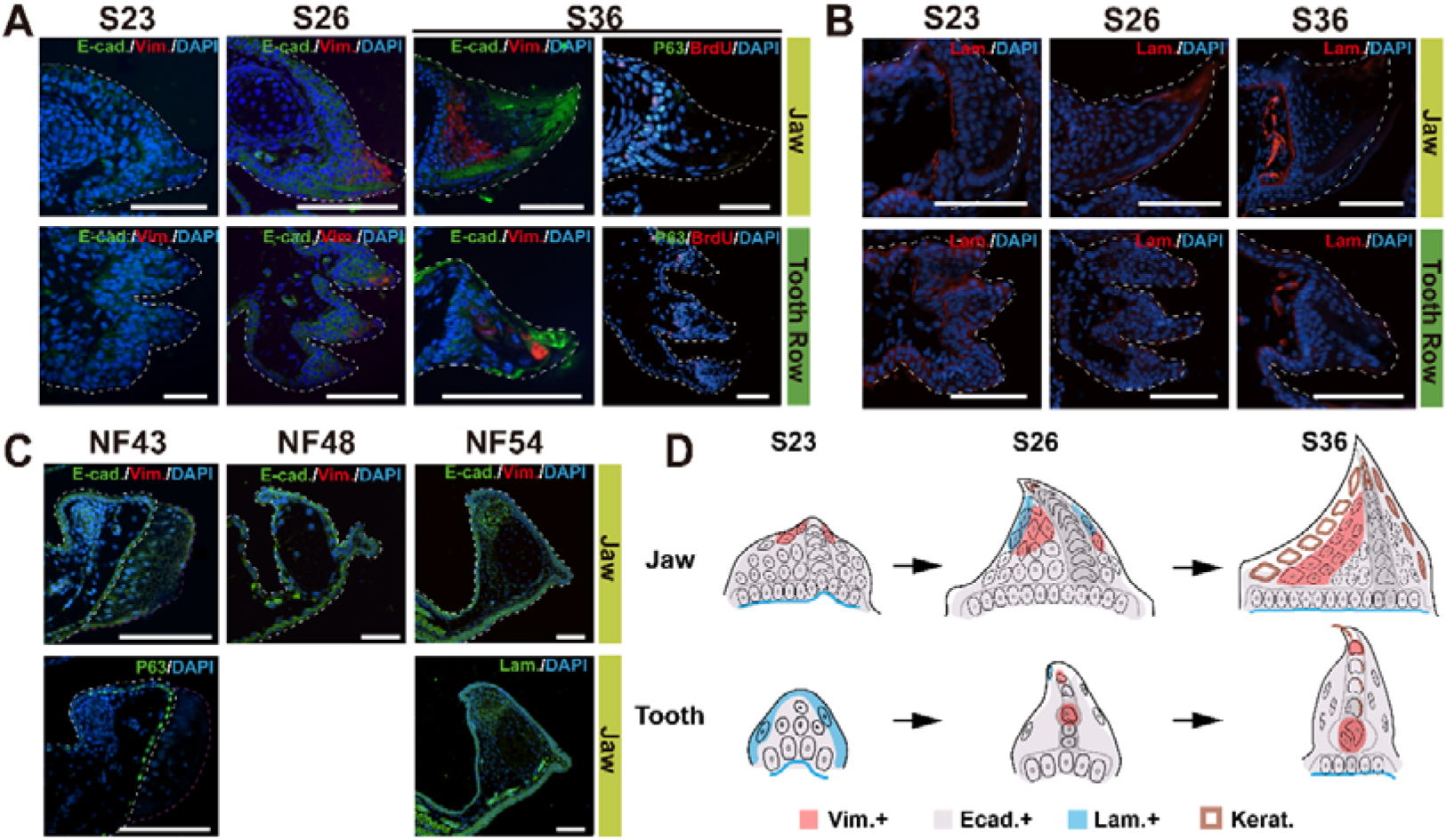
Spatiotemporal organization of oral epithelium during oral morphogenesis in *B. gargarizans* and *X. laevis*. A, Against a broadly E-cadherin-positive epithelial background, *vim* transcripts were first detected at the distal tips of the developing jaw and tooth ridges in S23 *B. gargarizans* tadpoles, before overt keratinization of these regions. Vimentin protein subsequently accumulated in suprabasal epithelial cells adjacent to the basal layer of the jaw and was excluded from the shingle-like cone-cell layer, in which E-cadherin remained enriched. In the tooth rows, vimentin was distributed in discontinuous distal and proximal domains associated with keratinized tooth structures, whereas E-cadherin remained broadly expressed throughout the differentiating epithelium. At S36, p63-positive basal cells and BrdU-labelled proliferating cells were detected in both the jaw and tooth epithelia. Vimentin-positive cells were not observed beneath the oral epithelial basement membrane in either region. Dashed lines delineate the boundaries of the oral epithelium. **Scale bars= 100**μ**m.** **B,** Throughout most stages of oral-disc morphogenesis of *B. gargarizans* tadpoles, laminin delineated the basal epithelial interface. At S26, basal laminin staining was markedly reduced, whereas during active jaw-sheath formation an additional laminin-positive domain appeared within the superficial keratinizing epithelium. Thus, oral-disc morphogenesis generally retains a laminin-positive basal boundary while undergoing transient matrix-associated remodeling within the superficial keratinizing layer. Dashed lines delineate the boundaries of the oral epithelium. **Scale bars= 100**μ**m.** **C,** Vimentin was undetectable in the oral epithelium of *X. laevis* tadpoles at NF43, NF48 and NF54, despite widespread E-cadherin expression. p63-positive basal epithelial cells were present at NF43, and laminin continued to delineate the basement membrane of the residual jaw epithelium at NF54. Dashed lines delineate the boundaries of the oral epithelium. **Scale bars = 100**μ**m.** **D,** Schematic summary of marker dynamics during jaw and tooth morphogenesis from S23 to S36, showing the emergence of regionally distinct epithelial states in the jaw and labial tooth primordia. In the jaw, the early E-cadherin-positive oral epithelium is underlain by a laminin-positive basal boundary, followed by the progressive emergence of vimentin-positive epithelial cells and keratinized elements as the jaw elongates and differentiates. At S26, laminin staining along the basal boundary is reduced in both the jaw and tooth primordia. By S36, the jaw comprises a basal epithelial layer, an expanded population of vimentin-and E-cadherin-double-positive inner epithelial cells, and keratinized sheath-cell and cone-cell derivatives. In the tooth primordium, laminin is initially present along the basal boundary and distal margin, followed by the appearance of spatially restricted vimentin-positive epithelial states and the progressive formation of keratinized tooth cusps. **Abbreviations: E-cad:** E-cadherin; **Vim:** vimentin; **Lam:** laminin; **Kerat:** keratinized structures.

In *X. laevis*, epithelial elaboration remained limited after mouth opening (Fig. 1J). Two epithelial folds, one directed upward and the other outward, formed over the lower jaw but remained only one to two cells thick and lacked keratinization, intraepithelial vimentin-positive domains, and differentiated cell types equivalent to those observed in *B. gargarizans* (Figs. 1H, 2C, S3). No corresponding epithelial fold developed over the upper jaw (Fig. 1G). Transient protrusions along the lower lip subsequently regressed during the developmental interval corresponding to oral-disc regression in metamorphosing *B. gargarizans* tadpoles (Fig. 1J; Fig. S3). Thus, *X. laevis* initiated oral epithelial outgrowth but did not execute the regionalized epithelial thickening, cell-state transitions and keratinization that generate the jaw, labial tooth rows and persistent papillae characteristic of *B. gargarizans*.

**Fig. 3.**
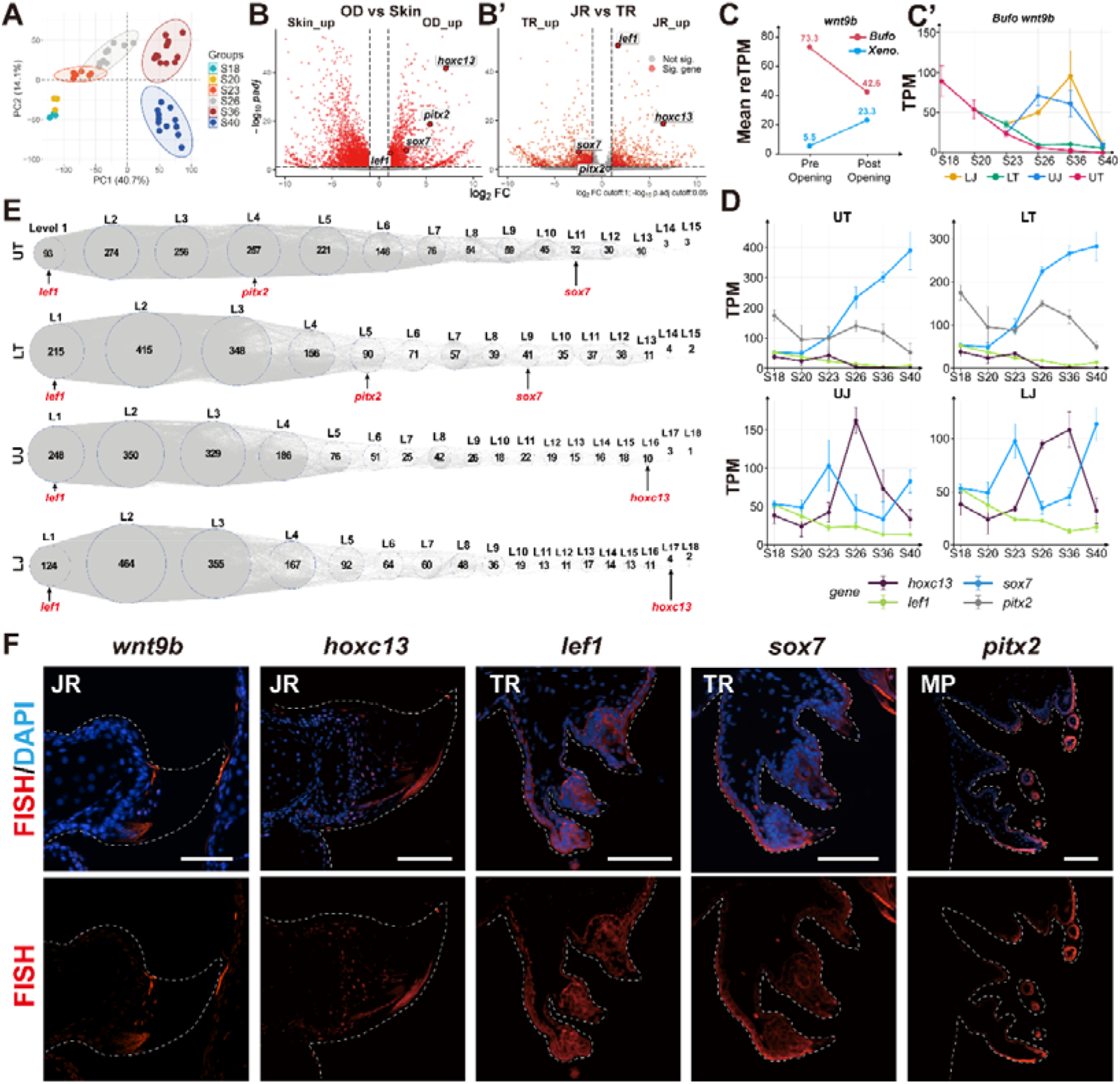
**Stage-and region-specific molecular programs during oral-disc morphogenesis in *Bufo gargarizans* tadpoles A,** Principal component analysis (PCA) of transcriptomes from oral-disc epithelial tissues sampled across developmental stages between S18 and S40. Each dot represents a biological replicate, with colors denoting developmental stage. Samples separated primarily according to developmental stage, demonstrating that developmental progression is the major source of transcriptional variation accompanying oral-disc formation, maturation and regression. **B–B**′, Volcano plots showing differentially expressed genes (DEGs) at S36 between the **OD** and ventral skin **(B)** and between pooled **JR** and pooled **TR (B**′). Red dots indicate significantly differentially expressed genes, and grey dots indicate genes that did not meet the significance thresholds. Dashed lines indicate the cutoffs for differential expression. Compared with ventral skin, the oral disc exhibited elevated expression of candidate regulatory and signaling-associated genes, including *hoxc13, wnt9b, pitx2, yap1, sox7, fzd9, ctnnb1* and *lef1*. Genes upregulated in the oral disc were enriched for developmental regulatory processes and keratinized-structure differentiation. In the JR–TR comparison, *hoxc13, wnt9b, wnt4* and *fzd9* were preferentially expressed in the jaw regions, whereas *sox7* and *yap1* were enriched in the tooth-row regions. *ctnnb1, lef1* and *pitx2* showed only modest regional differences or were close to the significance thresholds, indicating limited transcriptional divergence between jaw and tooth-row regions for these genes. **C-C’,** Expression of *wnt9b* before and after primary mouth opening in *B. gargarizans* tadpoles (C), and across developmental stages (C′). **D,** Temporal expression dynamics of selected candidate regulators. Upper panels show expression of *hoxc13*, *lef1*, *sox7* and *pitx2* in pooled TR and JS from S26 to S40, whereas the middle and lower panels show their expression in individual UT, LT, UJ, and LJ from S18 to S40. Expression levels are shown as transcripts per million (TPM); points indicate means and error bars indicate standard deviations (SD). Within the pooled TR, *sox7* increased from the onset of keratinization through maturation and regression, whereas *pitx2* declined after S26; *lef1* and *hoxc13* remained at relatively low levels throughout this period. Within the pooled JR, *hoxc13* peaked at S26 before declining, whereas *sox7* increased at S40. Across individual subregions, *lef1* gradually decreased during development; *sox7* progressively increased in UT and LT and remained highly expressed at S36–S40; *pitx2* displayed stage-dependent dynamics within the tooth-row regions; and *hoxc13* peaked during jaw differentiation and keratinization, at S26 in UJ and S36 in LJ. **E,** Time-ordered gene co-expression network (TO-GCN) analysis of transcription factors in the four oral-disc regions (UT, LT, UJ and LJ). Circles represent sequential virtual co-expression levels, with the number of transcription factors assigned to each level shown inside each circle. Grey lines indicate inferred co-expression relationships. Candidate transcription factors are highlighted in red beneath the corresponding levels, indicating their relative positions within the inferred developmental sequence. Across all four regions, the majority of transcription factors were assigned to early network levels, suggesting widespread transcriptional activation during early oral-disc development. *lef1* was consistently assigned to the earliest level in all four regions, supporting a conserved early regulatory role across oral-disc structures. In contrast, *pitx2* and *sox7* occupied later levels in the tooth-row networks, whereas *hoxc13* was assigned to the terminal levels of the jaw-region networks, indicating region-specific temporal deployment of candidate regulatory programs. Mfuzz clustering further resolved dynamic transcriptional programs associated with oral-disc development. **F,** HCR RNA–FISH analysis of selected regulators at approximately S36. Upper panels show merged FISH and DAPI signals, and lower panels show FISH signals alone. *hoxc13* was enriched in the keratinizing jaw sheaths, particularly within the shingle-like cone-cell layer. *lef1* and *sox7* were localized to the tooth-row epithelium; *lef1* was broadly distributed throughout the ridge epithelium, whereas *sox7* was concentrated in the outer epithelial layer of the most peripheral tooth ridge and was additionally detected in the keratinized jaw sheaths. *pitx2* was enriched in the marginal papillae (MP), particularly in the outer epithelium of the papillary projections. Dashed lines delineate the analyzed oral-disc structures. **Scale bars=100**μ**m**. **Abbreviations: JR,** jaw regions; **LM,** lower mandible; **LJ,** lower jaw sheath; **LT,** lower tooth row; **MP,** marginal papillae; **OD,** oral disc; **TR,** tooth-row regions; **UJ,** upper jaw sheath; **UT,** upper tooth row.

Together, these findings indicate that oral epithelial morphogenesis diverges following mouth opening, producing distinct trajectories of oral-disc elaboration in the two species. Histological analysis across larval development detected neither a cellular dermis nor localized mesenchymal condensations beneath the oral epithelium in either species. In *B. gargarizans*, the spatially ordered jaws, tooth-rows and papillary architectures therefore emerged within a continuous epithelial sheet (Fig. 2D). These observations identify the epithelium as the proximate basis of tadpole oral-disc patterning and promote the question of which spatially organized molecular programs specify individual oral modules.

### Molecular programs pattern the tadpole oral disc

Comparison of orthology-matched transcriptomes before and after mouth opening revealed divergent developmental programs in the oral tissues of *B. gargarizans* and *X. laevis* (Figs. 3, S4-8). Before mouth opening, *wnt9b* was expressed at substantially higher level in *B. gargarizans* than in *X. laevis* (Fig. 3C). In *B. gargarizans*, *wnt9b* expression declined during mouth opening and subsequently became enriched in the jaw epithelium, while remaining undetectable in the underlying cartilage and subepithelial tissues (Fig. 3C, C′, F). These findings implicate support a role for epithelial wnt signaling in oral-disc patterning before mouth opening.

Transcriptomic profiling of anatomically defined oral-disc epithelia revealed pronounced temporal and regional molecular patterning during oral-disc development in *B. gargarizans* (Figs. 3, S9-11). Across developmental stages, oral-disc epithelia occupied distinct transcriptional states associated with specification, maturation and regression (Fig. 3A; Fig. S11A-D). Within stage-matched samples, the upper and lower jaw regions showed closely related expression profiles, as did the corresponding upper and lower tooth-row regions (Fig. 3D; Fig. S8). These similarities suggest that paired jaw and tooth-row regions represent reiterated molecular modules within a common oral epithelial field.

The regulatory landscape identified by spatially resolved profiling comprised an early, broadly deployed oral-disc program and later module-specific differentiation programs (Figs. 3D-F, S9-S11). Genes enriched in oral-disc tissues included regulatory and signaling factors implicated in developmental patterning and keratinized epithelial differentiation (Figs. S9-11). Among the candidate regulators, *lef1* showed broad early enrichment and occupied at the earliest inferred tiers of temporally ordered co-expression networks in both jaw and tooth-row regions (Fig. 3E). In contrast, *sox7* was deployed later and was preferentially associated with tooth-row regions, whereas *hoxc13* was enriched in jaw regions during oral-disc differentiation (Fig. 3E). Oral-disc morphogenesis therefore proceeds from a shared early regulatory state towards regionally specialized programs that distinguish tooth rows from keratinized jaw structures.

Spatial expression analysis confirmed distinct expression domains for these candidate transcription factors within the oral epidermis (Figs. 3F, S2E). *lef1* transcripts were detected throughout the epithelia of the jaws and tooth rows, but were largely excluded from the keratinizing cone-cell layer of the jaws (Figs. 3F, S2E). *hoxc13* was enriched throughout the differentiating jaw epithelium, with expression extending from the basement layer to the keratinizing cone-cell layer (Figs. 3F, S2E), consistent with its association with jaw-specific epithelial differentiation (Fig. 3B′, D, E). *sox7* expression was restricted to the jaw tips, the labial epithelia of the outermost upper and lower tooth rows, and the precusp-cell domains of tooth ridges (Figs. 3F, S2E). In contrast, *pitx2* was enriched in the outer epithelial layer of the tooth ridges and papillae (Figs. 3F, S2E).

Collectively, these expression domains subdivide the oral disc into molecularly distinct epithelial territories corresponding to the jaw, tooth row and marginal papilla modules, thereby linking histological modularity to a temporally ordered and spatially specified molecular program (Fig. 3D-F). The epithelial restriction of *wnt9b*(*10*), together with the spatial distributions of other transcription factors, supports a model in which oral-disc initiation and modular patterning are driven predominantly by epithelium-intrinsic programs.

### Functional perturbation resolves distinct epithelial modules in the oral disc

CRISPR-Cas9 induced indel mutagenesis of *lef1*, *hoxc13*, *sox7*, and *pitx2* revealed that distinct oral-disc structures are governed by separable epithelial-enriched regulatory programs (Figs. 4A-F, S12-13).

**Fig. 4.**
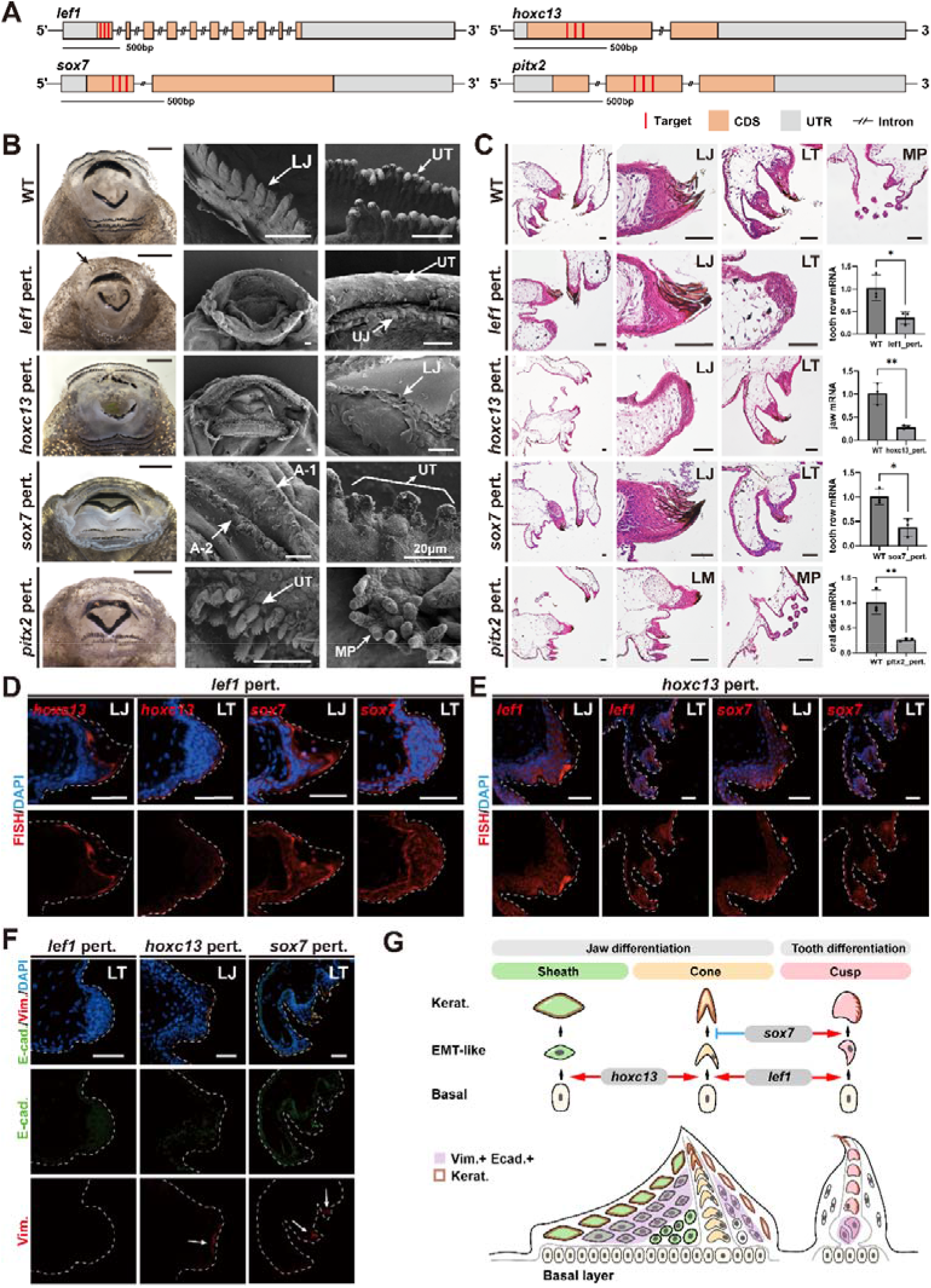
Functional perturbation of selected regulators reveals modular genetic control of tadpole oral-disc morphogenesis in *B. gargarizans* **A,** CRISPR/Cas9 targeting strategy for *lef1*, *hoxc13*, *sox7* and *pitx2*. Red lines indicate sgRNA target sites; apricot boxes indicate coding sequences (CDS); grey boxes indicate untranslated regions (UTRs); double slashes indicate omitted intronic intervals. **B,** Gross morphology and scanning electron micrographs of wild-type and gene-perturbed oral discs. Wild-type tadpoles develop organized keratinized jaw sheaths and labial tooth rows. Perturbation of *lef1* almost completely abolishes labial tooth rows (black arrow); *hoxc13* selectively disrupts keratinized jaw sheaths; *sox7* produces shortened and disorganized labial teeth while largely preserving jaw sheaths; and *pitx2* disrupts tooth-row organization and enlarges marginal papillae with clustered surface protrusions. **Scale bar=500**μ**m (gross views) and 50**μ**m (SEM),** unless otherwise indicated. **C,** Histological characterization and qPCR validation of gene perturbations. *lef1* perturbation reduces the tooth-row field to a poorly organized epithelial fold lacking terminal keratinized cusps. *hoxc13* perturbation abolishes characteristic jaw cone-and sheath-cell architecture, leaving only distal keratinized remnants. *sox7* perturbation preserves overall jaw organization but produces thinner tooth ridges with fewer internal cells and markedly reduced jaw cartilage. *pitx2* perturbation disrupts sheath-cell organization, tooth stratification and keratinization, and induces hypertrophic cauliflower-like marginal papillae. qPCR confirms reduced expression of each targeted gene relative to wild type. Bar plots show mean ± SD; \**P* < 0.05, \*\**P* < 0.01. **Scale bars=50**μ**m.** **D,** HCR RNA-FISH analysis of *lef1*-perturbed oral discs. *hoxc13* remains associated with residual keratinizing jaw epithelium but is restricted to the outer keratinizing surface of residual tooth ridges, whereas *sox7* is maintained in basal epithelial domains of both jaw and tooth-row regions. Dashed lines delineate the boundaries of the oral epithelium. **Scale bars=50**μ**m.** **E,** HCR RNA-FISH analysis of *hoxc13*-perturbed oral discs. *lef1* and *sox7* remain expressed in both disrupted jaw and retained tooth-row epithelia. Dashed lines delineate the boundaries of the oral epithelium. **Scale bars=50**μ**m.** **F,** Immunofluorescence staining for E-cadherin and vimentin in gene-perturbed oral discs. *lef1* perturbation markedly reduces E-cadherin and vimentin in the tooth-row region. *hoxc13* perturbation reduces jaw E-cadherin while residual vimentin-positive cells persist adjacent to disrupted keratinized tissue. *sox7* perturbation preserves basal E-cadherin but causes opposite regional changes in vimentin, reducing its signal in tooth ridges while redistributing it toward the jaw cone-cell layer. White arrows indicate residual vimentin-positive cells. Dashed lines delineate the boundaries of the oral epithelium. **Scale bars=50**μ**m.** **G,** Proposed model of epithelial-state transitions underlying modular oral-disc differentiation. Basal epithelial cells progress through EMT-like intermediate states toward module-specific terminal keratinized cell types in jaw and tooth structures. Perturbation analyses support distinct regulatory functions, with *hoxc13* promoting jaw epithelial differentiation and keratinization, *lef1* maintaining basal epithelial organization and tooth-row formation, and *sox7* exerting opposite effects on EMT-like differentiation in jaw and tooth modules. The lower schematic integrates these trajectories within the mature oral-disc architecture, highlighting a continuous basal layer, vimentin-positive/E-cadherin-positive internal epithelial populations and spatially restricted terminal keratinized derivatives. Brown regions labelled “Kerat.” indicate terminally keratinized cells. **Schematics are not to scale.** **Abbreviations: CDS,** coding sequence; **E-cad,** E-cadherin; **LJ,** lower jaw; **LM,** lower mouthpart; **LT,** lower tooth row; **MP,** marginal papillae; **UJ,** upper jaw; **UT,** upper tooth row; **UTR,** untranslated region; **Vim,** vimentin; **WT,** wild type.

Although *lef1* was broadly expressed in both jaw and tooth-row epithelia during early oral-disc development, its perturbation most severely affected tooth-row formation (Fig. 4B-C). In *lef1*-perturbed tadpoles (Fig. S12A), E-cadherin staining was nearly undetectable across the oral disc, and vimentin-positive cells were absent from the presumptive tooth-row region (Fig. 4F; Fig. S13B), indicating widespread disruption of epithelial cohesion and organization, together with failed tooth-row differentiation (84/96). Correspondingly, the tooth epithelium organized into a single shallow epithelial fold without cusp cells keratinization (Fig. 4B-C). Jaw elements remained recognizable but were shortened, less keratinized and showed reduced cone-cell differentiation (Fig. 4B-C). These findings identify *lef1* as a broad regulator of oral-disc epithelial architecture, with an especially strong requirement for the establishment and differentiation of labial tooth-row (Fig. 4G).

*hoxc13* perturbation preferentially impaired jaw morphogenesis while largely sparing tooth-row patterning (Fig. 4B-C). In *hoxc13*-perturbed tadpoles (Fig. S13B), jaw epithelial proliferation was reduced, and cone-cell and sheath-cell differentiation was impaired (96/100), yielding in a thinner epithelium (Figs. 4B-C, S13A). An epithelial basement layer remained histologically discernible despite the loss of p63 activity (Fig. S13A). The residual jaw epithelium failed to generate the cone cells and sheath cells required to construct the keratinized jaw sheath (Fig. 4C). Consequently, both the upper and lower jaw sheaths were severely disrupted, and the continuous keratinized sheath and serrated cutting edge were replaced by discontinuous keratinized remnants (Fig. 4B-C). E-cadherin staining was difficult to detect in the jaws (Figs. 4F, S13B), whereas tooth-row organization, distal cusp differentiation, and the typical 2(2)/3 tooth-row formula were largely retained (Fig. 4B). These data identify *hoxc13* as a predominantly jaw-specific regulator required primarily for cone-cell and sheath-cell differentiation and demonstrate that jaw keratinization is developmentally separable from that of the tooth-ridge (Fig. 4G).

In *sox7*-perturbed tadpoles (Fig. 12C), overall oral-disc organization and keratinized jaws were retained, although the jaw and tooth-row regions exhibited divergent responses (Fig. 4B-C). The jaw epithelium expanded, with a thickened basement layer and increased epithelial growth, whereas cone-cell and sheath-cell differentiation remained largely unaffected (Fig. 4C). By contrast, tooth ridges displayed an outwardly arched epithelial basal layer, reduced cellular density and impaired differentiation into fully keratinized tooth cusps (Fig. 4C). Consistent with these defects, terminal cusps and stratified tooth series were malformed, producing shortened, sparse and poorly organized labial teeth with reduced keratinization and disrupted tooth-row continuity (74/80). Vimentin signal was reduced in tooth ridges but redistributed within the jaw, shifting from the wild-type sheath-cell domain toward the cone-cell layer in perturbed tadpoles (Figs. 4F, S13B). Jaw-supporting cartilage was also markedly reduced despite expansion of the keratinized jaw epithelium (Fig. 4C), indicating that *sox7* perturbation uncouples epithelial jaw epithelial expansion from the formation of its supporting cartilage. Together, these finding suggest that *sox7* coordinates jaw size with subsequent labial tooth growth and differentiation (Fig. 4G).

Following *sox7* perturbation, β-catenin signaling was reduced in the basal jaw epithelium, and *lef1* and *hoxc1*3 expression were markedly diminished in this domain (Fig. S13C, G). Conversely, terminal cone-cell keratinization was enhanced (Fig. 4C). The vimentin-positive domain normally located proximal to the sheath-cell layer was lost, whereas E-cadherin and vimentin were ectopically co-expressed in cone cells (Fig. 4F), a pattern absent from wild-type tadpoles.

*Pitx2* perturbation strongly disrupted the peripheral papillae and reduced jaw sheath differentiation (Fig. 4B-C). In *pitx2*-perturbed tadpoles (Fig. S12D), tooth ridges contained fewer internal cells, failed to generate properly stratified tooth series and showed reduced keratinization (Fig. 4C). Jaw cone-cell morphology remained largely intact, whereas sheath-cell organization was disrupted and keratinization was reduced on both the lingual and labial surfaces. The most pronounced phenotype occurred in the marginal papillae (Fig. 4B-C), which became hypertrophic and cauliflower-like, resembling mucus glands morphologically(*11*). Overall, *pitx2* perturbation produced an enlarged oral disc with hypertrophic marginal papillae and defective labial tooth rows (63/80). These phenotypes indicate that *pitx2* constrains oral-disc growth and is required for the normal morphogenesis and patterning of marginal papillae.

Together, these perturbation experiments resolve the tadpole oral disc into partially independent epithelial modules controlled by distinct programs governing cellular adhesion, epithelial basement identity, cellular proliferation, cytoskeletal remodeling, and keratinization (Fig. S14). Perturbation of epithelial-restricted regulators selectively collapsed, expanded, or mispatterned specific structures, supporting a model in which the tadpole oral disc develops as a self-organized epidermal organ assembled from functional modules (Fig. 4G). These findings support a model in which epithelial-restricted molecular signaling programs are sufficient to pattern the distinct functional modules of the tadpole oral disc in the absence of a discernible canonical mesenchymal signaling center.

## Discussion

### Regionalization of oral-disc epithelial modules

Perturbation of selected epithelial-enriched transcription factors showed that the tadpole oral disc is patterned by separable regulatory programs controlling epithelial architecture, basal-cell identity, proliferation, cytoskeletal state transitions and keratinized differentiation (Fig. 4B-G, Fig. S14). These results define a regionally organized regulatory architecture in which shared epithelial programs are redeployed to generate distinct morphogenetic and differentiation outcomes in the jaw, labial tooth rows, and marginal papillae (Fig. 4G).

Among the factors examined, *lef1* emerges as the broadest upstream regulator, with context-dependent developmental outputs in the tooth-row and jaw epithelia. In the tooth-row field, *lef1* downregulation maintains the epithelium basement an undifferentiated state and prevents normal tooth-ridge formation, thereby compromising tooth-row folding and the keratinization tooth cusps (Fig. 4C). These phenotypes support a two-step model in which *lef1* first enables the tooth-field epithelium to form ridges that define tooth-row number, length and position, and subsequently promotes cusp-cell differentiation within these folds. This early requirement raises the possibility that interspecific variation in tooth-row number is mediated, at least in part, by modulation of *lef1*-dependent patterning. In the jaw, *lef1* downregulation disrupts basal epithelial organization and cone-cell maturation, whereas sheath-associated keratinization is retained (Fig. 4B, C; Fig. S13A, B), indicating that jaw identity does not depend entirely on the labial tooth-row differentiation program. Thus, *lef1* is a major regulator of tooth-row epithelial patterning and jaw cone-cell maturation (Fig. 4B-D, F, G), and these findings are against a model in which the jaw is assembled as a serial array of labial tooth-like units.

*hoxc13* defines a jaw-specific epithelial differentiation program: its perturbation reduces epithelial proliferation, compromises cone-cell and sheath-cell differentiation, and disrupts jaw keratinization (Fig. 4B, C), consistent with its expression across basal and distal keratinizing epithelial cells (Fig. 3F). This requirement parallels the established roles of *hoxc13* in mammalian hair follicles and amphibian claws, where it has been implicated in keratinocyte differentiation and cornification(*12*). Notably, *lef1* perturbation markedly reduces jaw *hoxc13* expression but does not abolish sheath-cell-associated keratinization (Fig. 4D; Fig. S13E). Terminal jaw keratinization can therefore persist despite reduced hoxc13 expression, suggesting that it is supported by additional regulatory inputs. Together, *lef1* and *hoxc13* define two interconnected epithelial programs: a shared *lef1*-dependent program that patterns tooth-row folds, tooth cusps and jaw cone cells, and a jaw-enriched, *hoxc13*-associated program that couples basal epithelial proliferation with cone-and sheath-cell differentiation and terminal keratinization (Fig. 4G; Fig. S15).

Although the shared requirement for *lef1* suggests that jaw cone cells and tooth precusp cells engage a common upstream epithelial program, several observations indicate that these populations subsequently follow distinct differentiation trajectories (Fig. 4G). Vimentin-positive cells occupy different positions in jaw and tooth epithelia: they are excluded from the maturing cone-cell layer but remain associated with differentiating precusp cells (Fig. 2A, D). Consistent with this distinction, *sox7* perturbation enhances cone-cell keratinization in the jaws but severely impairs precusp-cell differentiation and cusp formation in the tooth rows, accompanied by a redistribution of vimentin-positive epithelial domains (Fig. 4 C, E; Fig. S13B). Jaw cone cells and tooth cusps therefore share a *lef1*-associated upstream program yet diverge in their dependence on *sox7*-sensitive epithelial state transitions during differentiation (Fig. 4G).

The oral disc enlargement and papillary hypertrophy caused by *pitx2* perturbation identify its role as a constraint on peripheral epithelial expansion and a coordinator of organ-scale growth (Fig. 4B-C). Its intermediate position within the TOGCN coincides with the developmental emergence of peripheral oral-disc structures (Fig. 3E), supporting a model in which *pitx2* links organ-wide growth control to the subsequent patterning of non-keratinized marginal structures (Fig. S2E). By allowing jaw, tooth-row and papillary traits to vary semi-independently, this modular regulatory architecture could generate selectable phenotypic combinations, thereby providing a developmental substrate for oral-disc diversification.

### The tadpole oral disc as a transient, self-organizing epidermal organ

A notable feature of oral-disc morphogenesis in *Bufo* tadpoles is the absence of an identifiable dermal condensation or a morphologically distinct mesenchymal layer beneath the developing oral epithelium (Figs. 1C, D, I, 2A). No comparable subepithelial mesenchymal layer has been reported in other anuran larvae, including *Xenopus* and *Rana*(*6, 13*) (Fig. 5B). This organization distinguishes the oral disc from canonical vertebrate epithelial appendages(*14, 15*) and suggests that regional patterning and morphogenesis can proceed without a morphologically discrete mesenchymal organizer.

**Fig. 5.**
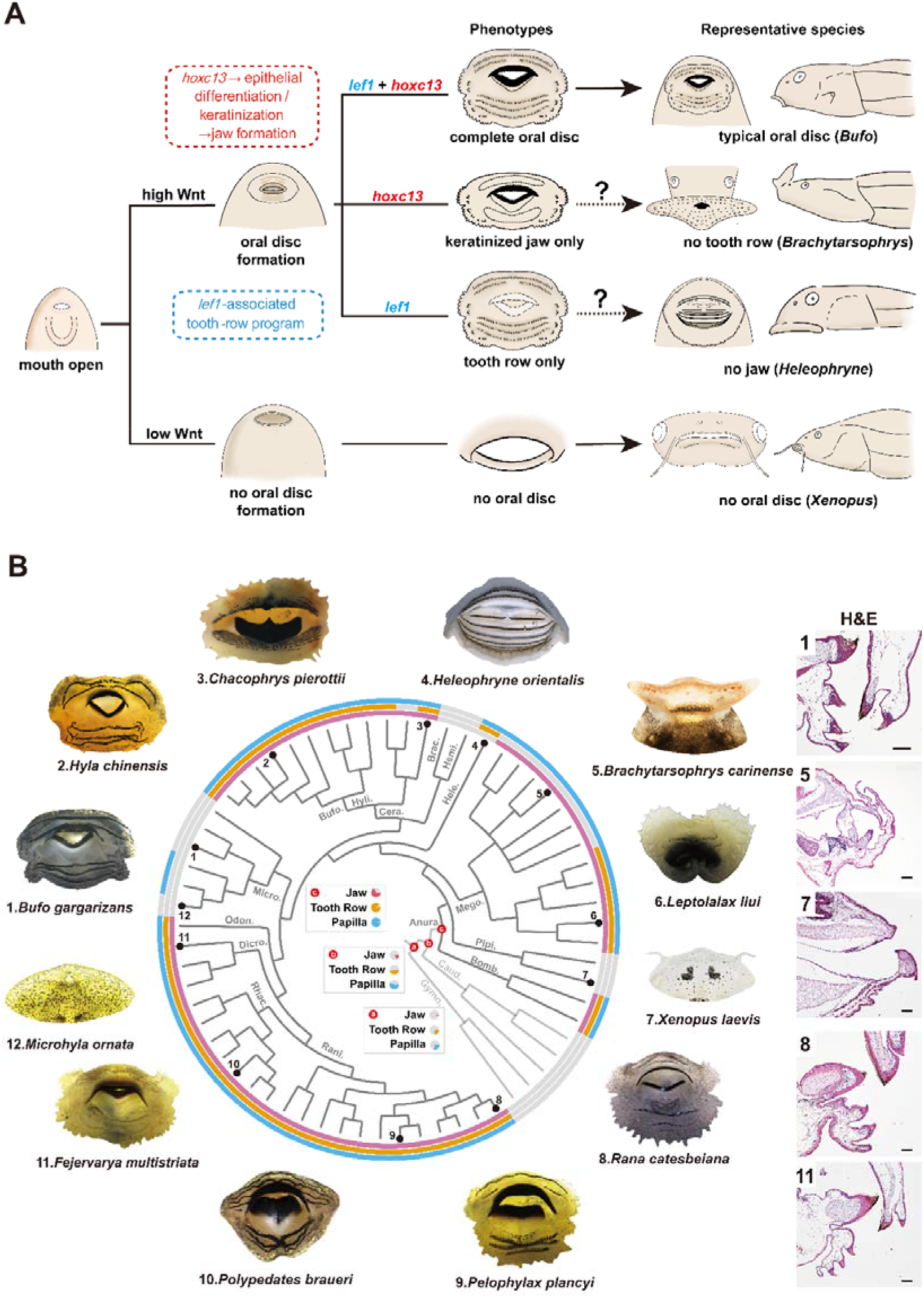
Modular assembly and evolutionary diversification of tadpole oral-disc architectures across amphibians A, Proposed developmental model for the modular diversification of tadpole oral morphologies. The model proposes that anuran larval oral diversity arises through differential activation, assembly, retention, reduction or loss of epithelial modules within the tadpole oral apparatus. Following mouth opening, high *Wnt* activity promotes oral-disc formation, whereas reduced *Wnt* activity is associated with failure or loss of oral-disc formation, as exemplified by *Xenopus*, which lacks a typical keratinized oral disc. In taxa with a complete oral disc, a *Lef1*-associated tooth-row program and an *Hoxc13*-dependent jaw-sheath differentiation and keratinization program are assembled to generate the keratinized feeding apparatus exemplified by *Bufo*. Selective deployment of individual modules produces alternative oral configurations, including a jaw-sheath-dominant state lacking tooth rows, represented by *Brachytarsophrys*^38^, and a putative tooth-row-dominant state lacking jaw sheaths, represented by *Heleophryne*^39^. The question mark denotes a predicted module combination that has yet to be functionally validated. Overall, the model suggests that repeated gains, reductions and losses of the oral disc can be explained by differential combinations of separable keratinized epithelial programs rather than by modification of the oral disc as an indivisible structure. **B,** Phylogenetic distribution of tadpole oral structures and representative epithelial architectures across amphibians. Circular phylogeny summarizing the distribution of major larval oral traits across sampled amphibians, including representative anuran lineages and caudate and gymnophionan outgroups. Tip-associated characters summarize the presence or absence of keratinized jaw, labial tooth rows and marginal papillae. Representative gross oral morphologies surrounding the tree illustrate the diversity of sampled oral configurations, including complete oral discs, jaw-or tooth-row-reduced states, oral-funnel-bearing forms and taxa lacking a typical oral disc. Numbered images correspond to the numbered tips in the phylogeny and to the H&E-stained sections shown at right. Histological sections highlight differences in epithelial organization, keratinized jaw-and tooth-row architecture, and the reduction or loss of oral-disc-associated structures among representative taxa. Gross images are not to scale. Pie charts labelled a–c at the center of the phylogeny depict the marginal probabilities of alternative ancestral states for each oral structure. **Scale bars=50**μ**m** for the H&E images. Species sampling and character-scoring criteria are provided in Table S1, respectively; ancestral-state reconstructions are shown in Fig. S17.

Vimentin and E-cadherin are widely used as markers of mesenchymal and epithelial identity, respectively. In the tadpole oral disc, however, vimentin-positive cells were restricted to the jaw and tooth-row epithelia, and no vimentin-positive subepithelial tissue was detected (Figs. 2A, S2B-C). This distribution is consistent with a specialized epithelial state and does not support the presence of a conventional subepithelial mesenchymal organizer(*16*). Vimentin can be induced in keratinocytes during migration, wound repair and differentiation-associated remodeling without loss of epithelial identity(*17*); its expression alone therefore is insufficient to infer mesenchymal lineage identity(*18*). Moreover, vimentin and E-cadherin exhibited extensive spatiotemporal overlap and coordinated redistribution during oral-disc development, most prominently between S26 and S36 (Figs. 2A, S2B-C). By S36, sheath cells became progressively spindle-shaped along the basal-to-distal differentiation axis, concomitant with reduced E-cadherin and increasing vimentin expression; neither marker was detected in terminally keratinized, anucleate cells. Comparable co-expression states have been described during development and regeneration as partial epithelial-to-mesenchymal transition (EMT) or hybrid epithelial-mesenchymal states(*16, 19*), in which epithelial cohesion is retained while cellular plasticity and remodeling capacity are enhanced. In regenerating epidermis, keratinocytes at migratory fronts can transiently express vimentin alongside keratins, and vimentin-dependent cytoskeletal remodeling contributes to keratinocyte migration and re-epithelialization(*17, 18*). Oral-disc morphogenesis may therefore transiently engage an EMT-like program that facilitate remodeling without compromising tissue continuity.

The onset of vimentin expression after precone-cell specification, together with its loss during metamorphic regression, is consistent with a transient role in epithelial differentiation and argues against a persistent signaling domain (Figs. 2A, S2C). Vimentin was also absent from the maturing cone-cell layer, further restricting its expression to an intermediate phase of epithelial remodeling. These temporal dynamics provide little support for assigning the vimentin-positive cells a dermal or mesenchymal signaling role(*16*).

Laminin provides complementary evidence that oral-disc remodeling occurs within an epithelial architecture organized around an intact basement membrane (Figs. 2B, D, S2D). The transient recruitment of laminin into the superficial keratinizing epithelium during jaw-sheath formation may establish a local matrix scaffold that supports assembly of the terminal keratinized structure while preserving the underlying basal matrix (Figs. 2B, D, S2D). In zebrafish fins, laminin-α5-associate epidermal cells at the distal tip have been proposed to coordinate fin-fold outgrowth and maintenance through extracellular signaling or structural support(*20*). A comparable mechanism may operate in the tadpole jaw, where localized matrix deposition could stabilize rapidly remodeling apical epithelial domains. Laminin enrichment in distal or terminal epidermal regions of developing fish fins and *Bufo* digits (Fig. S2A) further raises the possibility that this matrix-associated mechanism is recurrently deployed during the formation of rapidly remodeling epidermal structures.

The functional uncoupling of vimentin-positive domains organization from terminal jaw differentiation further constrains the likely role of this EMT-like epithelial state. The continued differentiation of cone cells after displacement of the normal vimentin-positive domain argues against an instructive mesenchyme-like function for these cells (Figs. 4F, S12B). Following *sox7* perturbation, the altered spatial relationship between vimentin-positive and E-cadherin-positive domains instead indicates a redistribution of epithelial differentiation states across the jaw epithelium. The EMT-like program therefore appears to promote local epithelial plasticity and tissue remodeling while preserving overall epithelial cohesion. Together, these findings support a model in which transient, regionally regulated epithelial-state transitions drive the formation of a complex vertebrate feeding apparatus without an identifiable mesenchymal organizer.

### Rapid oral disc assembly generates selectable variation

Comparative anatomy reveals no clear homologue of the oral disc in other vertebrate larvae. Salamander (Fig. S16) and caecilian larvae lack an equivalent integrated feeding apparatus(*21*), whereas superficially similar keratinized structures in other aquatic vertebrates (e.g., lampreys(*22*)) probably evolved convergently. Fish unculi, for example, comprise individual keratinized epidermal cells and lack the multicellular organization of the tadpole oral disc(*23*). The lamprey sucking disc likewise develops only after metamorphosis(*22*), contrasting with the larval origin of the anuran oral disc.

The anatomical discontinuity between tadpoles and adults reflects the extensive developmental reorganization imposed by anuran metamorphosis(*24*). By decoupling larval and adult feeding ecologies, metamorphosis relaxes developmental and functional constraints that would otherwise couple their morphologies, allowing the feeding apparatus at each stage to respond independently to distinct functional and selective demands(*25, 26*). Closely related species with relatively conserved adult morphologies can therefore differ markedly in tadpole oral-disc architecture(*27*). The oral disc represents a larva-specific epithelial innovation that has contributed to the ecological diversification of anuran tadpoles(*24, 26, 28*).

Functionally, the tadpole oral disc forms a transient yet mechanically resilient interface that enables tadpoles to exploit diverse freshwater resources, including algae, periphyton, detritus and suspended particles(*7, 28, 29*). Continuous local keratinization rapidly replenishes surfaces abraded during feeding(*30*), resembling the sustained growth of avian beaks(*31*) and mammalian nails(*32*) but differing from the periodical renewal of scales, hair, feathers and teeth(*4, 14, 15*). Canonical epithelial appendages typically rely on reciprocal epithelial-mesenchymal signaling to maintain organ identity and progenitor activity over prolonged periods(*4, 14, 15*). By contrast, the tadpole oral disc uses an epidermis-centered program adapted for rapid larval assembly, continuous repair and dismantling at metamorphosis(*7, 33*). This strategy combines mechanical resilience with the developmental plasticity required for rapid remodeling(*34*).

Such plasticity is evident in the diversification of oral-disc architectures across anurans (Fig. 5)(*7, 9, 28, 35*). Typical pond-type tadpoles, including those of *Bufo*, *Hyla*, *Rana*, *Pelophylax*, *Polypedates* and *Fejervarya*, retain the canonical combination of keratinized jaw, labial tooth rows and marginal papillae(*7, 36, 37*) (Fig. 5B). Ancestral-state reconstruction suggests that these modules were established separately and subsequently underwent lineage-specific retention, modification, reduction or loss (Figs. 5B, S17B-D). Our perturbation experiments support this modular organization by showing that individual components can vary semi-independently, producing combinations resembling those of extant anurans. *lef1* perturbation eliminated labial tooth rows while retaining prominent jaws, resembling the jaw-dominant condition of *Brachytarsophrys* (Fig. 5B)(*38*). Conversely, *hoxc13* perturbation disrupted jaw sheaths while largely preserving tooth rows, resembling the jaw-reduced, tooth-row-rich condition of *Heleophryne* (Fig. 5B)(*39*). The distinct effects of *sox7* and *pitx2* further indicate partially independent control of jaw size, tooth differentiation and papilla morphogenesis. Concurrent attenuation of multiple epithelial programs produces simplified oral fields resembling those of *Xenopus* (Figs. 1F, 5A, B) and microhylid tadpoles(*7, 26*).

These experimental phenotypes do not establish that the corresponding natural morphologies must have evolved through changes in the genes examined here. Instead, they demonstrate that differential modulation of shared epithelial programs is sufficient to generate structural combinations already represented in extant species. Variation in the timing, magnitude and spatial deployment of these programs could therefore produce diverse oral phenotypes from a common developmental toolkit (Fig. 5A). Variation in oral-disc position and orientation, together with differences in the number, shape and arrangement of jaw sheaths, tooth rows and papillae, further expands oral-disc morphospace(*7, 8, 28*). How the number, morphology and spatial arrangement of individual labial tooth rows are specified remains unresolved(*9*).

Together, these findings provide a developmental framework for the repeated modification, rearrangement and loss of oral-disc modules across anuran phylogeny (Figs. 5, S17). This developmentally accessible variation provides a substrate for natural selection, enabling alternative combinations of oral-disc modules to be assembled and refined under different ecological conditions. Such combinations support feeding modes including suction, grasping, scraping and filter feeding(*7, 29*).

## Conclusion

Together, our findings establish the tadpole oral disc as a self-organizing epidermal system that develops without a discernible subepithelial mesenchymal signaling center. Spatially restricted epithelial regulatory programs specify and differentiate distinct oral regions, enabling jaw sheaths, labial tooth rows and marginal papillae to develop as partially autonomous modules. Perturbing *lef1*, *hoxc13*, *sox7*, and *pitx2* produced module-specific phenotypes resembling combinations that have evolved repeatedly across anurans, linking differential deployment of shared epithelial programs to selectable variation in larval feeding structures. Thus, epithelial self-organization enables a transient organ to combine rapid assembly and repair with structural complexity and evolutionary flexibility.

## Materials and Methods

### Phylogenetic backbone

The phylogenetic framework for the selected taxa was conducted based on recent studies of amphibian phylogeny(*40*). Sixty-four representative species spanning diverse larval oral morphologies were selected, and their phylogenetic relationships were inferred using the Phylogeny Subsets Tool of VertLife (https://vertlife.org/phylosubsets/). A total of 1,000 phylogenetic trees were generated, from which a consensus tree was inferred using the R package phytools. A complete species list together with the corresponding morphological characters are provided in Table S1. Tree editing, annotation, and visualization were performed using TreeViewer v2.2.0(*41*) and Adobe Illustrator.

### Ancestral state reconstruction

Ancestral state reconstruction of oral disc characters was performed in *R* using the *ace* function in the *ape* package. The consensus tree was first converted into a fully dichotomous tree using *multi2di*, and oral disc characters were coded as discrete traits. Three discrete character evolution models were evaluated: equal rates (ER), all rates different (ARD), and symmetrical rates (SYM). Marginal ancestral states were reconstructed under each model, and model fit was assessed using log-likelihood values. The ER model exhibited the highest log-likelihood and was therefore selected for the final analyses. The reconstructed marginal likelihoods for each internal node under the ER model are provided in Table S2.

### Animal rearing

Adult *Bufo gargarizans* were purchased from a branch of Nanjing Toad Farm, and *Xenopus laevis* were obtained from Minhang Laboratory Animal Center, East China Normal University. Natural mating was performed following the protocol of Lane & Khokha(*42*). Embryos and tadpoles were maintained in a controlled aquatic facility at Minhang Laboratory Animal Center at 20∼25°C under veterinary supervision. All experimental procedures were approved by East China Normal University (Approval No. ARXM2024010) and were conducted in accordance with institutional guidelines.

### Staging and sampling

Developmental staging of *Bufo gargarizans* followed established staging system(*43*), which comprises 46 developmental stages spanning fertilization to the completion of metamorphosis(*43*). Six representative stages were selected to encompass the major morphogenetic events during *Bufo gargarizans* tadpole mouthpart development, corresponding to Gosner stages S18, S20, S23, S26, S36, and S40 (*43*). S18 corresponds to the initial invagination of the stomodeum, marking the onset of stomodeum formation(*44*); S20 represents perforation of oral-membrane; S23 marks the initiation of oral disc development before overt keratinization; S26 corresponds to the establishment of the basic oral structures; S36 represents completion of oral disc morphogenesis; and S40 marks the onset of oral disc degenration. For *Xenopus laevis*, which does not develop a typical oral disc(*45*), developmental stages were assigned according to the Nieuwkoop and Faber (NF) staging system(*46*).

Samples was collected at NF26, NF32, NF43, NF48, NF54, and NF58, corresponding approximately to the selected developmental stages of *B. gargarizans* based on the comparative developmental framework of McDiarmid and Altig(*7*). Additional wild tadpole specimens used in this study were obtained from the Vertebrate Collection of Biological Museum, East China Norma University and were not deposited with catalog numbers.

To assess cell proliferation, BrdU (1 mg/mL, Sigma® B5002) was administered by injection 2∼5 hours before sampling. Tadpoles were euthanized using 0.1% eugenol (1:1,000 dilution). Tissues for RNA extraction were dissected and immediately placed in RNAlater (Invitrogen®, AM7020). Tissues collected for genotyping were placed in 75% ethanol immediately after dissection. Tissues for histological staining and *in situ* hybridization (ISH) were fixed in 4% paraformaldehyde for 2 hours, rinsed in phosphate-buffered saline (PBS), and dehydrated through a graded ethanol series (30%∼ 100%).

### Histological sections and staining

Mouthpart samples were embedded in paraffin and sectioned sagittally at a thickness of 7μm using a Leica RM2235 microtome. Sections were floated on a 44°C water bath and processed for hematoxylin and eosin (H&E) staining(*47*), ISH, and immunofluorescence (IF) staining. BrdU assays were performed following the protocol of Lee *et al*.(*48*), using an anti-BrdU antibody (Servicebio®, GB12051). For IF staining, sections were deparaffinized in xylene, rehydrated through a graded ethanol series, and subjected to antigen retrieval in 1× sodium citrate buffer at 95°C for 12 min. Sections were subsequently digested with 20 µg/mL proteinase K (NEB® P8107S) for 1min at room temperature, blocked with 3% bovine serum albumin (BSA) for 30 min at room temperature, and incubated with primary antibodies overnight at 4 °C.

Primary and secondary antibodies used for IF staining included anti-E-cadherin (Invitrogen®, PA5-85088), anti-vimentin (Invitrogen®, MA5-11883), anti-laminin (Sigma-Aldrich®, L9393), anti-p63 (Servicebio®, GB150065), anti-β-catenin (Servicebio®, GB150016). The corresponding secondary antibodies were goat anti-rabbit IgG Alexa Fluor 488 (Invitrogen®, R37116), goat anti-rabbit IgG Alexa Fluor 594 (Invitrogen®, R37117), goat anti-mouse IgG Alexa Fluor 488 (Invitrogen®, R37120), and goat anti-mouse IgG Alexa Fluor 594 (Invitrogen®, R37121). Following three washes in PBS, sections were incubated with fluorescent secondary antibodies and counterstained with DAPI. Fluorescence images were acquired using a Nikon Ni-E microscope equipped with a Digital Sight 10 camera.

### RNA extraction and quantitative RT-PCR

Total RNA was extracted using TRIzol (Invitrogen, 15596018) following the manufacturer’s instruction. RNA quality and concentration were assessed using a NanoDrop spectrophotometer (Thermo Fisher Scientific). First-strand cDNA was synthesized using HiScript III 1st Strand cDNA Synthesis Kit (Vazyme, R312).

qPCR was performed using Hieff® SYBR Green Master Mix (Yeasen®, 11202ES08) on an Applied Biosystems QuantStudio 5 Real-Time PCR system. Thermal cycling conditions consisted of an initial denaturation at 95°C for 5 minutes, followed by 40 cycles of 95°C for 10s and 60°C for 30s. A melting-curve analysis was subsequently performed to verify amplification specificity by gradually increasing the temperature from 60 °C to 95 °C while continuously monitoring fluorescence. The qPCR primer sequences are listed in Table S3. All reaction were performed in technical triplicate. Relative gene expression levels were normalized to *actb* and calculated using the 2^(-ΔΔCt) method(*49*). The corresponding raw Ct values are provided in Table S4. Data are presented as mean ± standard deviation (s.d.). Statistical significance was determined using an unpaired two-tailed Student’s t-test, and differences with *p*<0.05 were considered statistically significant.

### RNA-seq and transcriptomic analyses

Because the *Bufo gargarizans* mouthpart undergo progressive morphological differentiation during development, RNA-seq sampling strategy were adapted according to developmental stage. Before S23, the entire oral region was collected because regional morphological differentiation had not yet occurred. At S23, the upper and lower mouthparts were dissected separately, corresponding to the onset of oral regionalization. From S26 onward, further anatomical specialization enabled independent sampling of the upper tooth row (UT), lower tooth row (LT), upper jaw (UJ), and lower jaw (LJ).

To ensure sufficient RNA yield from tadpoles of different developmental sizes, stage-specific pooling strategies were adopted. Samples consisted of three individuals at S18, ten individuals per region at S26, and five individuals per region for all remaining stages. Three independent biological replicates were generated for each sample type (Table S5). All dissected tissues were immediately preserved in RNAlater before RNA extraction.

Total RNA extraction, cDNA library preparation, and sequencing were performed by Novogene Bioinformatics Technology Co., Ltd. Libraries were prepared using the NEBNext Ultra RNA Library Prep Kit for Illumina (NEB®, E7530L), and library quality was evaluated using an Agilent 5400 system. Qualified libraries were sequenced on the Illumina NovaSeq X Plus platform using a 150-bp paired-end (PE150) configuration, generating approximately 6 Gb of sequencing data per sample.

### RNA-seq data analysis

Raw sequencing reads were initially processed by Novogene® Bioinformatics Technology Co., Ltd. to remove adapter sequences. The resulting clean reads were further filtered using Trimmomatic v0.39(*50*) with the following parameters: LEADING:3, TRAILING:3, SLIDINGWINDOW:4:15, HEADCROP:10, and MINLEN:140. Filtered reads were aligned to the *Bufo gargarizans* reference genome (ASM1485885v1) using Hisat2 v2.2.1(*51*) with the -*dta* option to facilitate downstream transcript assembly. Alignments were sorted and converted to BAM format using SAMtools v1.10(*52*).

Transcript assembly and expression quantification were performed using StringTie v2.1.4(*53*) with the annotated *Bufo gargarizans* genome as the reference annotation(*54*). StringTie was executed with the *-e* option, and raw gene count matrices were extracted using the prepDE.py script. Gene expression levels were normalized as transcripts per kilobase million (TPM). Genes with a mean TPM ≤ 1 across all samples were excluded from downstream analyses. Differential expression analysis was conducted using DESeq2 (*55*) based on raw read counts, and functional enrichment analyses were carried out using clusterProfiler(*56*).

### Construction of the time-ordered gene co-expression network (TO-GCN)

Time-ordered gene co-expression network (TO-GCN) analysis was applied to infer the temporal ordering of transcription-factor (TF) expression, following previously established developmental transcriptomic workflows(*57–59*). Transcriptomic data from six developmental stages of *Bufo gargarizans* (S18, S20, S23, S26, S36, and S40) were used to construct TO-GCNs for four oral-disc subregions: upper tooth-row region (UT), upper jaw region (UJ), lower tooth-row region (LT), and lower jaw region (LJ). Only TF genes were included because they constitute the core regulatory components governing developmental gene-expression programs (Table S6).

For each subregion, TF genes were hierarchically organized into discrete TO-GCN levels according to their inferred temporal expression dynamics. Eighteen TO-GCN levels were identified for both UJ and LJ, whereas UT and LT comprised 11 and 15 levels, respectively. Lower TO-GCN levels contained TFs predominantly expressed during earlier developmental stages, whereas higher levels contained TFs with peak expression at later stages.

TO-GCN construction involved three major steps. First, Pearson correlation coefficients (PCCs) were calculated for all pairwise TF combinations, and significantly correlations (*p*<0.05) were retained. Second, gene co-expression networks were constructed from significant TF correlations. Third, TFs were assigned to sequential temporal layers using a breadth-first search (BFS) algorithm initiated from predefined seed genes(*57, 58*).

Because the oral disc remains morphologically differentiated at S18 and S20, transcriptomic data from these stages were treated as representing a common oral tissue across all subregions. From S23 onward, oral disc regionalization becomes apparent, although tooth row and jaw domains cannot yet be morphologically distinguished. Accordingly, transcriptomic data from S23 were subdivided into upper jaw and lower jaw datasets, which were subsequently used for UT/UJ and LT/LJ analyses, respectively.

TO-GCN construction followed the modified protocol of Wong et al.(*57, 58*), and all analysis scripts are publicly available at https://github.com/petitmingchang/TO-GCN_STAR-Protocol. Filtered TPM matrices were used as input, resulting in optimal PCC thresholds of 0.94 for UT, 0.94 for UJ, 0.93 for LT, and 0.94 for LJ (Supplementary Information). Initial seed genes were identified using the Monotonic Feature Selector(*60*) to detect monotonically decreasing expression patterns across developmental time. The highest-ranked TF genes in each subregion was selected as the initial seeds: E2F3 for UT, E2F3 for UJ, WDHD1 for LT, and SALL4 for LJ Supplementary Information. TFs were identified using AnimalTFDB v4.0(*61*) (https://guolab.wchscu.cn/AnimalTFDB4/#/TF_Predict), yielding 2,285 predicted TF genes in the *B. gargarizans* Supplementary Information.

### DIG-labeled RNA *in situ* hybridization (ISH)

DIG-labelled RNA in situ hybridization (ISH) was performed on paraffin-embedded sagittal sections of tadpole mouthparts to determine spatial gene-expression patterns. Antisense RNA probes targeting selected genes were synthesized, whereas corresponding sense probes served as negative controls.

Paraffin sections were deparaffinized in xylene, rehydrated through a graded ethanol series, and washed with PBST. Proteinase K digestion (5μg/mL) was performed for 10 min, followed by postfixation in 4% PFA. After dehydration and air-drying, hybridization buffer containing 50% formamide, 5× SSC, 0.1% Tween-20, and 0.6ng/μL DIG-labelled RNA probe was applied to each section. Slides were incubated at 65°C for 16 h.

Following hybridization, sections were washed three times with 2× SSC and subsequently with 0.2× SSC (20 min per wash at 65 °C) before rinsing in MABT. Sections were blocked in MABT containing Roche blocking reagent for 2 h, followed by incubation with anti-DIG alkaline phosphatase-conjugated antibody (1:300; Roche, 11093274910) overnight at 4 °C. Hybridization signals were visualized using NBT/BCIP alkaline phosphatase substrate (Roche, 11442074001), followed by PBS washes and image acquisition.

### Hybridization Chain Reaction (HCR)

The HCR RNA fluorescence in situ hybridization protocol for paraffin sections was adapted from the third-generation HCR v3.0 method and modified according to the Molecular Instruments protocol, while maintaining buffer compositions consistent with those reported previously(*62*). B3-series DNA hairpin amplifiers were used for signal amplification. Gene-specific probe sets were designed using the published HCR probe design pipeline(*63*).

### CRISPR-Cas9 functional validation

Target sgRNAs were designed using FlashFry(*64*). Candidate target sites were selected from the first coding exon downstream of the translation start codon, prioritizing sequences with high predicted on-target specificity and minimal off-target potential. Two to four sgRNAs were designed for each gene, and the corresponding target sequences, genome assembly versions, and GenBank accession numbers are provided in Table S3.

sgRNA templates were synthesized by *in vitro* transcription. Primer sequences are listed in Table S3. sgRNAs were synthesized using the MEGAscript T7 Kit (Invitrogen®, AM1334), purified by 7.5M LiCl precipitation, washed with chilled 75% ethanol, and validated by agarose gel electrophoresis.

Adult *B. gargarizans* were induced to undergo amplexus and ovulation by two injections of human chorionic gonadotropin (hCG). Fertilized eggs were collected, dejellied in 2% L-cysteine(*65*), and rinsed in 0.1× MMR. To assemble ribonucleoprotein (RNP) complexes, a injection mixture containing NLS-Cas9 nuclease, sgRNA, and 10× reaction buffer was prepared. Microinjection was performed at the one-cell stage, after which embryos were cultured in 0.1× MMR supplemented with 0.1% gentamicin.

### Genotyping and indel verification

To assess genome-editing efficiency, injected embryos were raised to the appropriate developmental stage, and tail tissues were collected for genotyping. Genomic DNA was extracted using a genomic DNA extraction kit (Tiangen®, DP304). PCR primers were designed to amplify genomic regions flanking each sgRNA target site. Editing efficiency was initially evaluated by Sanger sequencing of PCR amplicons, followed by targeted amplicon sequencing for high-resolution, allele-specific characterization of indel mutations. All primers used for genotyping and amplicon sequencing are listed in Table S3.

## Data availability

The RNA sequencing datasets generated in this study have been deposited in the NCBI Sequence Read Archive (SRA) under BioProject accession number PRJNA1493307 and will be made publicly available upon publication.

## Acknowledgements

S.W. acknowledges that the inspiration for this study originated from his work on edentulism during the ontogeny in 2016-2017. Over the following years, this observation gradually evolved into the present study. We thank all colleagues and collaborators for their valuable support and assistance throughout this research. We are especially grateful to M. Li for providing substantial assistance with the experiments carried out over the past field seasons and to X. Liao for assistance with paraffin sectioning. We sincerely thank Y. Zhang of East China Normal University and S. Qi of Sun Yat-Sen University for generously providing the specimens for comparisons. We also thank Z. Gong of East China Normal University and the East China Normal University Multi-Functional Platform for Innovation (004) for assistance with SEM imaging. S.W. acknowledges that this study received no funding and was completed at the researcher’s own expense.

## Author contributions

S.W. conceived the project. N.D. designed of the Bufo gene-editing experiments. C.C., N.D., J.H., and Y.S. performed the experiments and collected the data. C.C. and N.D. analyzed the data. S.W. and N.D. wrote the manuscript and developed the conceptual hypotheses. All authors read and approved the final manuscript.

