## Supplementary Information for "Epithelial Self-Organization Patterns the Tadpole Oral Disc"

1. **Developmental dynamics of the tadpole oral disc**

The tadpole oral disc of *Bufo gargarizans* develops through a highly ordered sequence in which an initially simple oral epithelium is progressively patterned into keratinized larval feeding structures, maintained as a functional oral apparatus, and subsequently dismantled during metamorphosis(*7, 28, 66, 67*). Across the stages examined here, oral-disc morphogenesis can be divided into four broad phases: pre-patterning, regional differentiation, functional maturation, and regression. These phases are reflected in coordinated changes in external morphology, epithelial histology, basement-membrane organization, proliferative activity, and the regional distribution of vimentin and E-cadherin (Figs. S1-2).

At S18, before overt oral-disc formation, the stomodeum is deeply invaginated and the oral membrane remains intact. Laminin outlines the basement membrane of the upper and lower oral epithelia and converges at the prospective mouth-opening site (Fig. S1A; Fig. S2D). By S20, as the primary mouth begins to form, the oral membrane is reduced to a thin residual epithelial partition. The laminin-positive basement membrane becomes locally attenuated and is subsequently breached at the prospective mouth-opening site, while the prospective jaw and tooth-row regions first become histologically recognizable as epithelial aggregates within the oral-disc field (Fig. S1A; Fig. S2D). At S23, epithelial thickening and deepening epithelial folds mark the onset of jaw- and tooth-row differentiation, and the supporting jaw cartilages begin to condense. During this transition, vim transcripts are first detected distally in the developing jaw and tooth ridges, whereas vimentin protein remains below detectable levels, indicating that transcriptional activation precedes overt vimentin protein accumulation. Laminin continues to delineate the basement membrane at this stage, suggesting that early epithelial folding proceeds along an intact basal matrix interface before overt keratinization (Fig. S1A; Fig. S2C-D).

By S26, black keratin first appears at the distal ends of the jaw sheaths and labial tooth rows, with keratinization initiating earlier and becoming more prominent in the jaws. BrdU labeling becomes largely restricted to the basal epithelial layer of the jaw and tooth-row ridges, indicating that proliferative activity is concentrated in basal and near-basal epithelial populations as terminal keratinization begins distally (Fig. S1A-B). At the same stage, vimentin protein is detected in the distal epithelium along emerging keratinization fronts. Laminin is reduced along parts of the basement membrane and becomes enriched in the outer keratinizing epithelium, revealing a redistribution of basement-membrane-associated matrix signals during epithelial differentiation and keratinized structure formation (Fig. S2B-C). By S36, the oral disc is fully mature and bears well-developed keratinized jaw sheaths and a 2(2)/3 labial tooth-row formula. Regression begins by S40, accompanied by reduced epithelial thickness, attenuation of the basal layer, and progressive withdrawal of keratinized structures (Fig. S1A, C). Oral-disc loss proceeds broadly in the reverse order of assembly, with jaw sheaths regressing before labial teeth (Fig. S1). By S42, the larval oral disc has been lost (Fig. S1A).

The dynamics of vimentin further resolve the epithelial transitions underlying oral-disc morphogenesis (Fig. S2C). At S20, vimentin-positive cells are absent from both the oral-disc epithelium and the underlying subepithelial region. By S23, vim transcripts are detected at the distal ends of developing jaw and tooth ridges by FISH, whereas vimentin protein remains undetectable by IHC. At S26, vimentin protein appears in the distal epithelium of both jaw and tooth ridges, coincident with the formation of keratinization fronts. At S36, jaw-associated vimentin is restricted to a triangular epithelial domain adjacent to the basal side of the jaw epithelium and is excluded from the shingle-like cone-cell layer. In tooth rows, vimentin occupies discontinuous distal and proximal epithelial domains associated with keratinized tooth structures. After regression at S42, the jaw-associated vimentin domain contracts and persists only beneath the upper jaw epithelium.

E-cadherin shows region-specific relationships with vimentin during oral-disc maturation and regression (Fig. S2B). In the jaws, E-cadherin-positive cells are broadly distributed within the oral epithelium at S26 and become concentrated in the unkeratinized shingle-like cone-cell layer by S36, with limited overlap with the vimentin-positive triangular domain. In tooth rows, E-cadherin extends broadly across the differentiating tooth ridge, including the vimentin-positive basal/proximal region. Thus, mature jaw and tooth-row epithelia exhibit distinct E-cadherin–vimentin configurations, consistent with region-specific epithelial states within the oral disc. During regression, E-cadherin becomes largely restricted to non-keratinized epithelial regions as keratinized oral structures are withdrawn.

p63 and BrdU staining further define the proliferative basal epithelial architecture of the mature oral disc (Fig. 2A; Fig. S1B). At S36, p63-positive cells form basal epithelial domains in both jaw and tooth-row regions, while BrdU-positive cells are detected in basal and near-basal epithelial regions. Together, these patterns indicate that the mature oral disc maintains regionally organized basal epithelial populations that support the keratinized jaw and tooth-row modules before their coordinated regression during metamorphosis. Combined with the vimentin and E-cadherin distributions, these data define a sequence of region-specific epithelial states that underlie the formation, functional maintenance, and metamorphic loss of the tadpole oral disc.

2. **Supplementary description of the functional perturbation results**

Additional histological and molecular analyses clarified how each transcription-factor perturbation reshaped epithelial organization in distinct regions of the tadpole oral disc (Fig. 4B-F; Fig. S12). In *lef1*-perturbed tadpoles, residual *lef1* expression was largely confined to tissue outside the oral-disc field. Within the oral disc, the broad reduction of E-cadherin coincided with loss of vimentin-positive cells from the presumptive tooth-row region, supporting a severe disruption of tooth-row epithelial identity(*16, 68*) (Fig. S12B). In the jaw, however, residual vimentin signal persisted along the inner margin of the keratinizing jaw tissue and extended into the region normally occupied by the shingle-like cone-cell layer in wild-type tadpoles (Fig. S12B). Because this cone-cell layer was absent or poorly organized after *lef1* perturbation, the altered vimentin distribution indicates that the residual vimentin-positive epithelium no longer retained its wild-type sheath-cell-associated organization (Fig. S12B). Histologically, the residual tooth-bearing region lacked a clear basal epithelial layer, contained loosely organized connective-tissue-like cells, and failed to generate terminal keratinized cusps (Fig. 4C). Occasional residual teeth were observed in some edited individuals, but organized tooth-row folding, stratified tooth formation, and mature cusp differentiation were largely absent. By contrast, jaw architecture was only partially affected: upper and lower jaw elements remained recognizable, the cone-cell layer was reduced, and lingual sheath-cell-associated keratinization was partly preserved (Fig. 4B). These observations indicate that *lef1* is especially required for tooth-field epithelial organization, tooth-row folding, and tooth-cusp differentiation, while the jaw sheath-cell keratinization program is comparatively less sensitive to *lef1* loss.

In *hoxc13*-perturbed tadpoles, H&E staining showed that a basal epithelial layer remained detectable in the disrupted jaw, indicating that loss of *hoxc13* did not completely eliminate jaw epithelial organization (Fig. 4C; Fig. S12A). However, this residual basal layer lacked the p63 activity characteristic of wild-type jaw basal cells, separating persistence of basal histological architecture from maintenance of basal-cell molecular identity (Fig. S12A). The remaining jaw epithelium was thin and poorly differentiated, yet still thicker than non-oral-disc epidermis, suggesting that *hoxc13* loss primarily impairs jaw-specific differentiation rather than epithelial survival (Fig. 4C; Fig. S12A). Consistent with this interpretation, *hoxc13* perturbation specifically blocked the differentiation of cone-cell and sheath-cell lineages that normally build the keratinized jaw sheath. The residual jaw epithelium resembled the poorly differentiated oral epithelium of *Xenopus* tadpoles, which lack typical keratinized jaw structures (Fig. 1F-H; Fig. S3). E-cadherin staining was weak or difficult to detect in the jaw but persisted in the tooth-row region, indicating partial preservation of tooth epithelial organization despite severe jaw defects (Fig. S12B). Vimentin signal was markedly reduced, with only faint staining near outer keratinizing cells of the upper and lower jaws and tooth ridges (Fig. S12B). Tooth ridges were moderately reduced, but distal cusps remained differentiated and keratinized, and the typical 2(2)/3 tooth-row formula was largely retained (Fig. 4B). Together, these data reinforce a regionally biased requirement for *hoxc13*, with the jaw sheath program showing strong dependence on *hoxc13* and the tooth-row program remaining comparatively preserved.

In *sox7*-perturbed tadpoles, residual *sox7* expression was concentrated in shingle-like cone cells of the jaws and in keratinizing tooth cusps (Fig. S12D). E-cadherin staining delineated the expanded jaw epithelium and the residual basal layer of tooth ridges, allowing vimentin redistribution to be mapped within defined epithelial regions (Fig. S12B). Vimentin showed opposite regional responses: it was weakened in tooth ridges but redistributed in the jaw, shifting from the wild-type sheath-cell-associated domain toward the cone-cell layer, where E-cadherin signal is normally reduced (Fig. S12B). Because *sox7* is expressed in the precusp-cell region, the tooth-row defects caused by *sox7* perturbation are consistent with impaired precusp-cell maturation and defective progression toward terminal cusp differentiation. Histologically, tooth ridges displayed outward bulging of the basal epithelium, reduced cellular density, and defective progression toward mature keratinized teeth. Terminal cusps were reduced or malformed, stratified tooth series failed to form properly, the first posterior labial tooth row frequently displayed a medial gap, and the posterior-most labial tooth rows persisted as ridge-like epithelial folds with reduced keratinization (Fig. 4C). In contrast, the jaw epithelium expanded, with a thickened basal layer and increased epithelial growth, while cone-cell and sheath-cell differentiation were largely retained (Fig. 4C). Jaw-supporting cartilages were markedly reduced despite expansion of the keratinized jaw epithelium, indicating that *sox7* perturbation uncouples epithelial jaw growth from development of the underlying skeletal support. These findings suggest that *sox7* influences both epithelial state positioning and tooth-row maturation, producing distinct, region-specific consequences in jaw and tooth-row epithelia.

In *pitx2*-perturbed tadpoles, *pitx2* expression was nearly undetectable (Fig. S12D), but basal organization and proliferative activity of the tooth-row epithelium were largely retained. Thus, *pitx2* loss did not broadly abolish the epithelial framework of the oral disc. In the jaw, cone-cell morphology remained largely intact, whereas sheath-cell organization was markedly disordered. Keratinization was weakened on the lingual side, where sheath cells normally appose the keratinized jaw tissue, and was also reduced on the labial side. In the tooth-row region, the basal epithelial layer arched outward, tooth ridges contained fewer internal cells, stratified tooth series failed to form normally, and keratinization was reduced despite retention of basal organization and proliferation (Fig. 4C). These changes indicate that *pitx2* acts after initial establishment of jaw and tooth-ridge epithelia to regulate epithelial growth, spatial organization, and keratinized differentiation. The most prominent phenotype occurred in marginal papillae, which underwent marked hypertrophy and formed clustered, cauliflower-like epithelial protrusions (Fig. 4B). These structures were morphologically reminiscent of mucus-gland-like outgrowths(*11*), although additional marker analyses would be required to establish glandular identity. Together, these supplementary observations indicate that *pitx2* primarily constrains peripheral oral-disc growth and papilla morphology, while also contributing to the ordered arrangement and keratinized differentiation of jaw and tooth-row structures.

Collectively, these perturbation analyses resolve the tadpole oral disc into regionally distinct epithelial programs with different transcription-factor dependencies. *lef1* perturbation preferentially disrupts tooth-field epithelial organization and cusp differentiation; *hoxc13* perturbation selectively compromises jaw sheath differentiation; *sox7* perturbation impairs tooth-row maturation while altering epithelial state positioning in the jaw; and *pitx2* perturbation most strongly affects papilla growth and the orderly differentiation of keratinized oral structures. These findings support a modular model in which jaw sheaths, labial tooth rows, and marginal papillae are generated by separable yet coordinated epithelial programs within the larval oral-disc field.

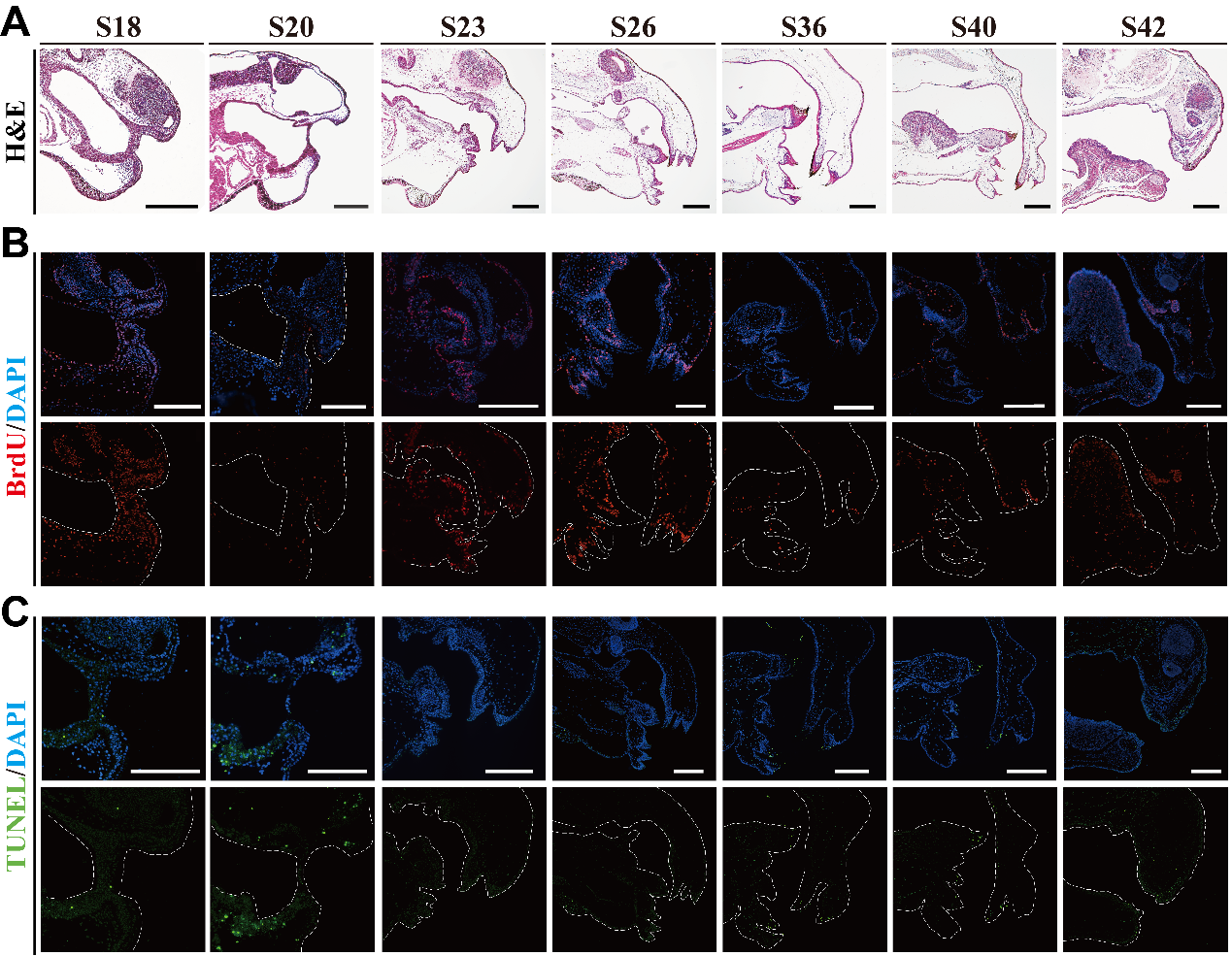

**Fig. S1 Full-field staining of developing tadpole oral structures in *Bufo gargarizans*.**

**A–C**, Full-field H&E, BrdU and TUNEL staining of the oral region across representative developmental stages. H&E staining shows progressive oral-disc morphogenesis from stomodeal invagination and oral-membrane thinning to jaw-sheath and tooth-row differentiation, maturation, metamorphosis and loss (**A**). BrdU staining shows that cell proliferation is broadly distributed in the early oral epithelium, becomes enriched in the basal epithelial layer of jaw sheaths and tooth rows during differentiation and maturation, and declines during metamorphosis (**B**). TUNEL staining shows that cell death is sparse at early stages and becomes more evident in the outer regions of jaw sheaths and tooth rows during metamorphic regression (**C**). **Scale bars = 250 μm.**

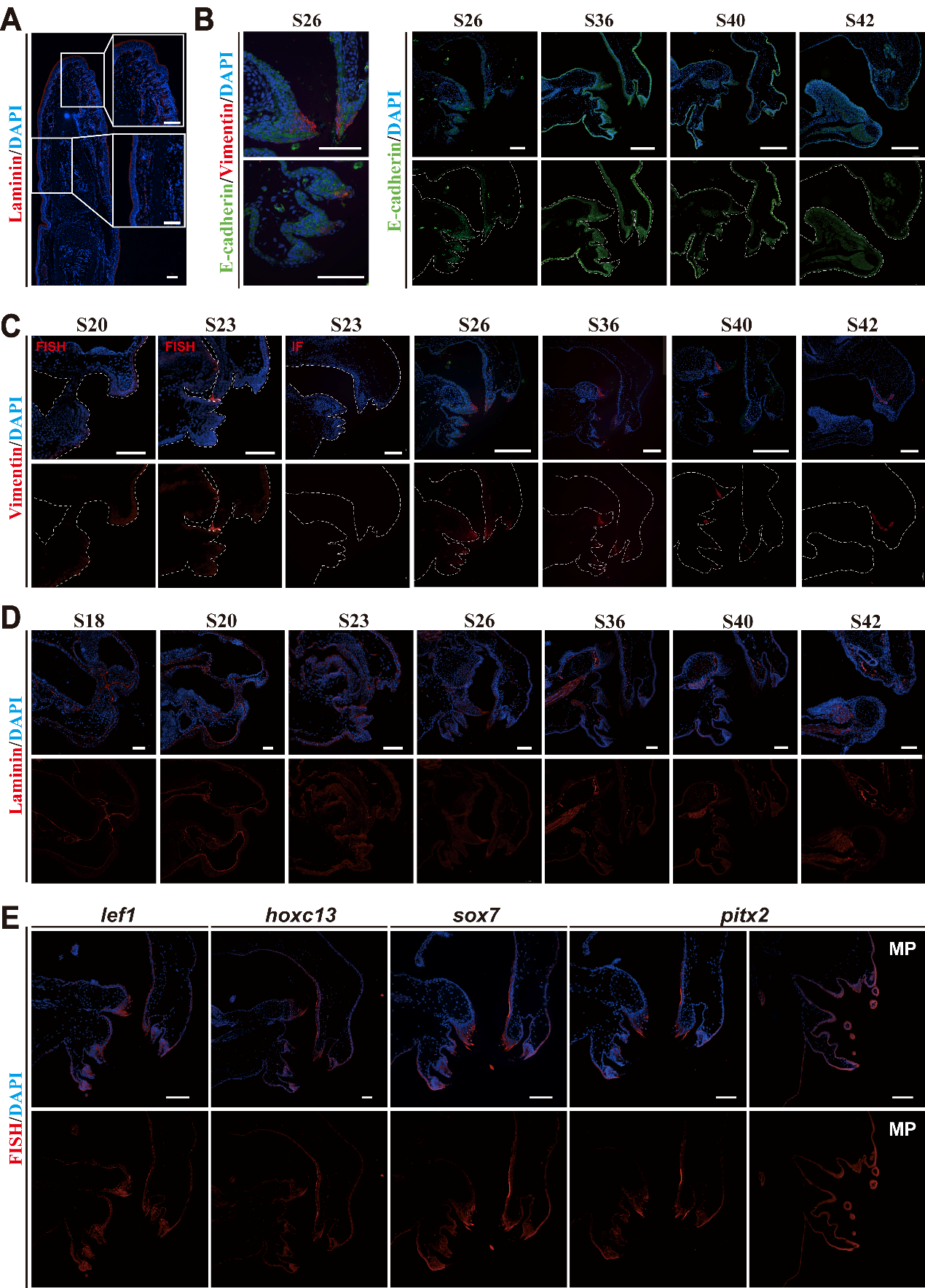

**Fig. S2 Full-field immunofluorescence and HCR RNA-FISH staining of developing tadpole oral structures in Bufo gargarizans.**

**A**, Laminin immunofluorescence staining of a Bufo digit, showing broad laminin enrichment along the outer epidermal surface. **B, Full-field E-cadherin immunofluorescence in the oral region at S26, S36, S40, and S42. E-cadherin is broadly detected throughout the oral epithelium, including the jaw-sheath and tooth-row territories, and remains evident during early metamorphic remodeling. The cone-cell layer is positive for E-cadherin but lacks detectable vimentin signal, whereas sheath cells show partial co-expression of E-cadherin and vimentin. In the tooth cusps undergoing keratinization, E-cadherin and vimentin signals substantially overlap.** **C**, Full-field vimentin staining in the oral region across representative developmental stages. HCR RNA-FISH shows that vim signal is absent or barely detectable at S20 and first appears at the distal tip of the developing jaw region at S23. Vimentin immunofluorescence from S23 to S42 shows that vimentin-positive cells localize mainly to the distal jaw region and tooth-row regions proximal to keratinized cusps, with signals retained in restricted epithelial regions during metamorphic remodeling. **D**, Full-field laminin immunofluorescence in the oral region from S18 to S42. Laminin outlines the basement membrane of the early oral epithelium, becomes locally reorganized during mouth opening and oral-disc differentiation, and is detected in basal and superficial epithelial-associated domains during jaw-sheath and tooth-row formation, maturation, and regression. **E**, Full-field HCR RNA-FISH images showing the spatial distribution of lef1, hoxc13, sox7, and pitx2 in wild-type tadpole oral discs at the stable stage. Upper panels show merged HCR and DAPI signals, and lower panels show HCR signals alone. lef1 signal is enriched in tooth-row epithelia, hoxc13 signal is enriched in the jaw-sheath region, sox7 signal is detected in tooth-row and adjacent epithelial regions, and pitx2 signal is detected in oral-disc epithelia and marginal papillae (MP). Dashed lines outline the oral-disc region. **Scale bars = 250 μm in A-C** **and 100 μm in D, E.**

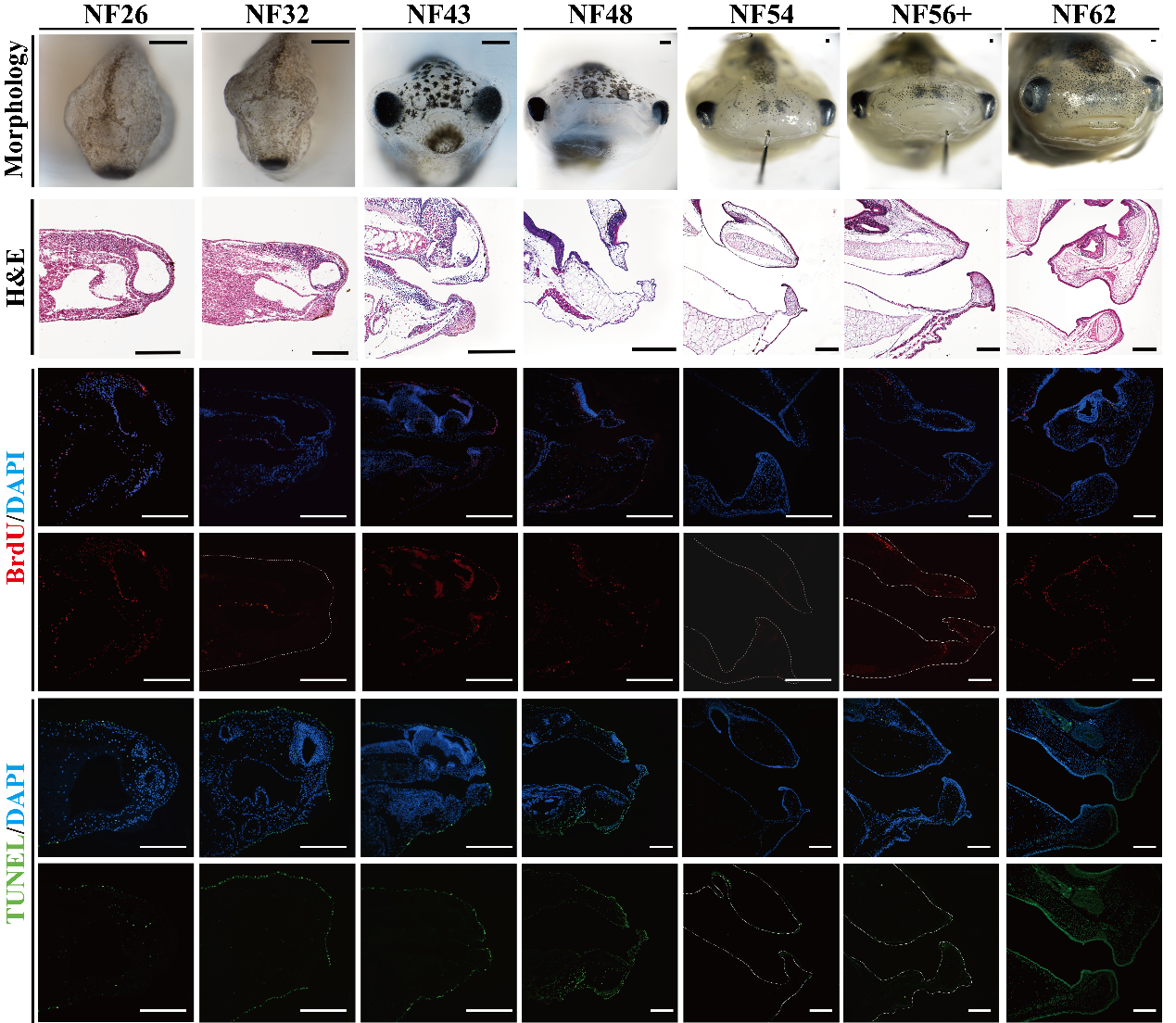

**Fig. S3 Developmental dynamics of tadpole mouthpart in *Xenopus laevis***

Gross morphology, histological sections, cell death, and cell proliferation were examined in the oral region of ***Xenopus laevis*** from NF26 to NF62. During early development, the presumptive mouth region and oral depression become discernible, but keratinized oral-disc structures are absent. As development proceeds, the oral margin expands and upper and lower jaw cartilages form, whereas typical anuran larval oral-disc structures, including keratinized jaw sheaths, labial tooth rows, and marginal papillae, are not detected. Cell proliferation is initially limited, becomes detectable in basal epithelial cells of the labial margin, and subsequently decreases, with signals mainly associated with intraoral cartilage or adjacent tissues and relatively weak proliferation in the oral-margin epithelium. Cell death is detected predominantly in the outer epidermis throughout development. **Scale bars = 250 μm.**

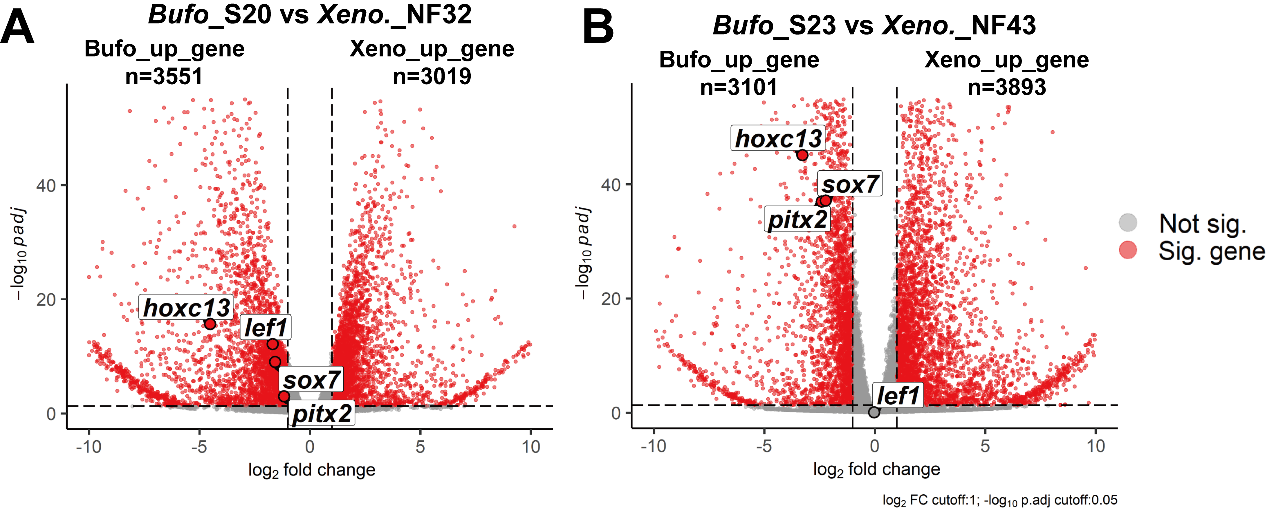

**Fig. S4 Cross-species transcriptomic comparisons around primary mouth opening in *Bufo* and *Xenopus*.**

Volcano plots showing differential expression of one-to-one orthologous genes between *Bufo gargarizans* oral samples and *Xenopus laevis* oral samples at two developmentally aligned stages before and after primary mouth opening. **A,** Comparison between *Bufo* S20 and *Xenopus* NF32, corresponding to the stage before primary mouth opening. A total of 3,551 orthologs showed higher expression in *Bufo*, and 3,019 orthologs showed higher expression in *Xenopus*. **B,** Comparison between *Bufo* S23 and *Xenopus* NF43, corresponding to the post-primary mouth-opening stage when oral-disc differentiation has begun in *Bufo*. A total of 3,101 orthologs showed higher expression in *Bufo*, and 3,893 orthologs showed higher expression in *Xenopus*. Red points indicate significantly differentially expressed orthologs, and dashed lines indicate the differential-expression thresholds. Selected oral-disc-associated regulatory and signaling genes are labeled. The *x* axis shows log_2_ fold change, and the *y* axis shows −log_10_ adjusted *P* value.

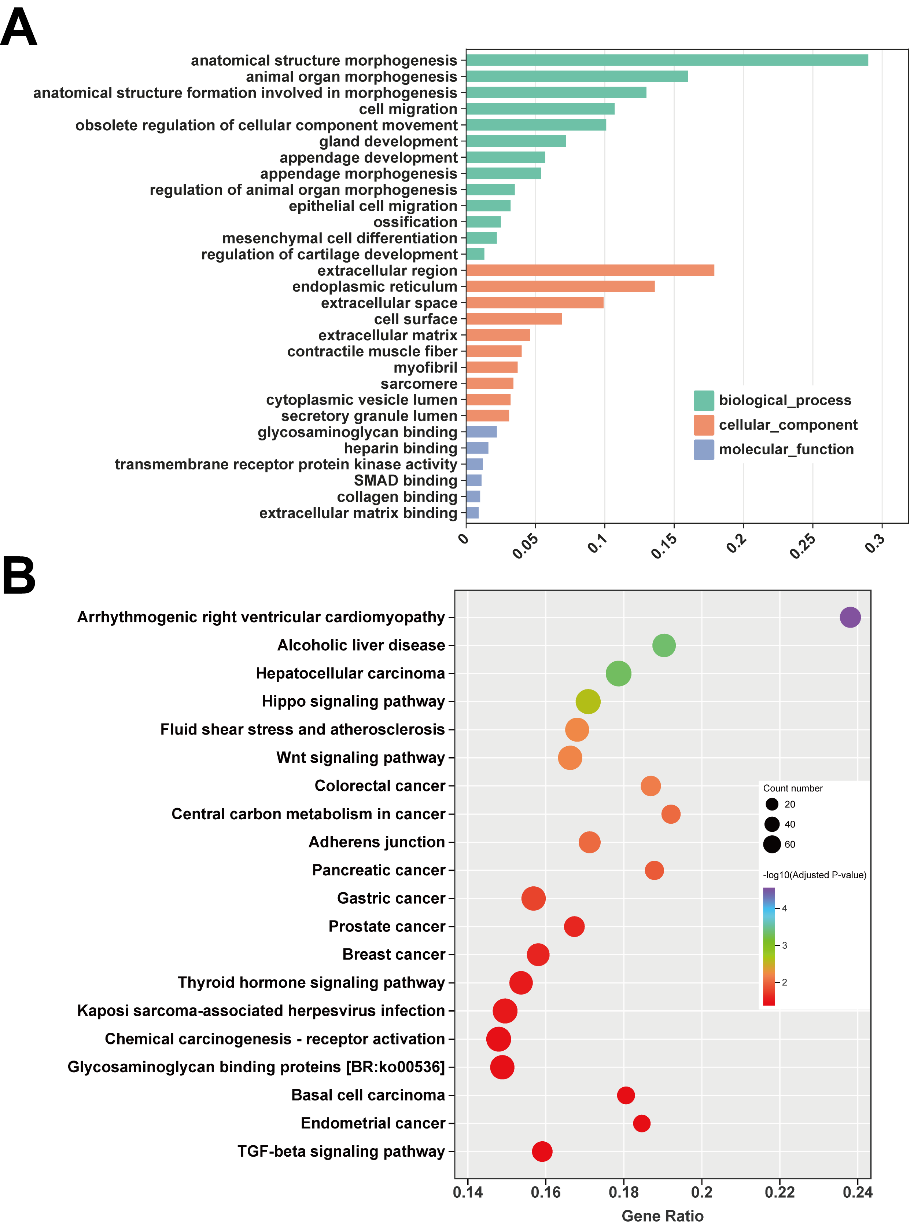

**Fig. S5 GO and KEGG enrichment analyses of *Bufo*-upregulated orthologs before primary mouth opening.**

**A,** GO enrichment analysis of one-to-one orthologous genes showing higher expression in *Bufo gargarizans* S20 oral samples than in *Xenopus laevis* NF32 oral samples. Enriched terms are grouped by biological process, cellular component, and molecular function. Bar length indicates GeneRatio.
**B,** KEGG enrichment analysis of the same *Bufo*-upregulated ortholog set. Bubble size indicates the number of genes assigned to each pathway, and color indicates −log_10_ adjusted *P* value.

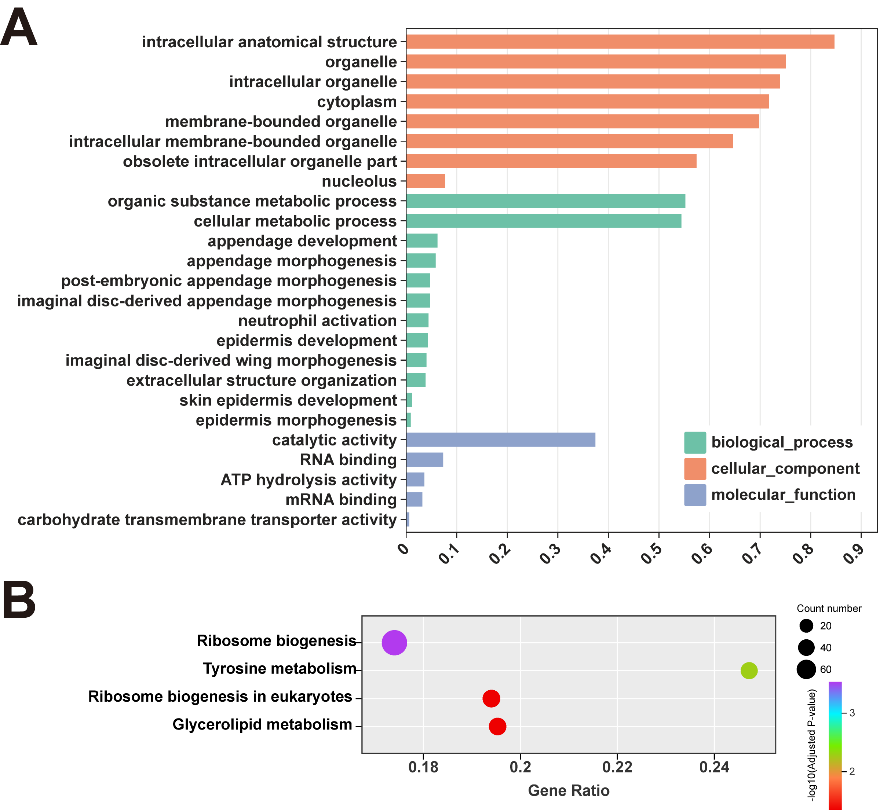

**Fig. S6 GO and KEGG enrichment analyses of *Bufo*-upregulated orthologs after primary mouth opening.**

**A,** GO enrichment analysis of one-to-one orthologous genes showing higher expression in *Bufo gargarizans* S23 oral samples than in *Xenopus laevis* NF43 oral samples. Enriched terms are grouped by biological process, cellular component, and molecular function. Bar length indicates GeneRatio.
**B,** KEGG enrichment analysis of the same *Bufo*-upregulated ortholog set. Bubble size indicates the number of genes assigned to each pathway, and color indicates −log_10_ adjusted *P* value. Representative enriched terms and pathways are shown.

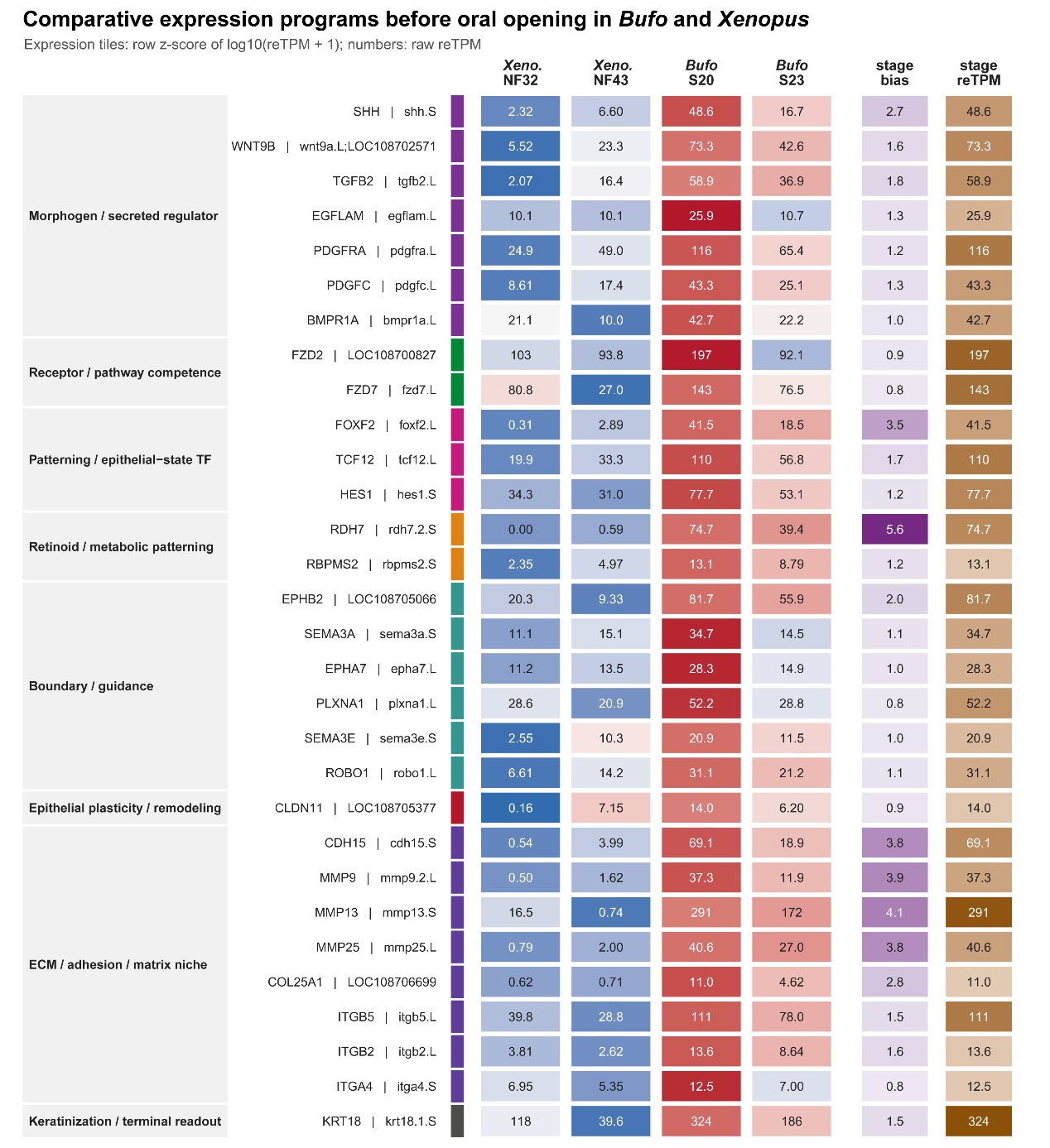

**Fig. S7 Comparative expression program before primary mouth opening in *Bufo* and *Xenopus*.**

Representative genes associated with the early oral-region program are grouped by functional category and compared across Xenopus NF32/NF43 and Bufo S20/S23. Tile colors indicate row z-scores of log10 (reTPM + 1), and numbers indicate raw reTPM values. The early program includes morphogen, receptor, patterning, boundary/guidance, epithelial-remodeling, ECM/adhesion and early keratinization-associated genes, suggesting that oral epithelial patterning and signaling competence are established before overt oral-disc differentiation.

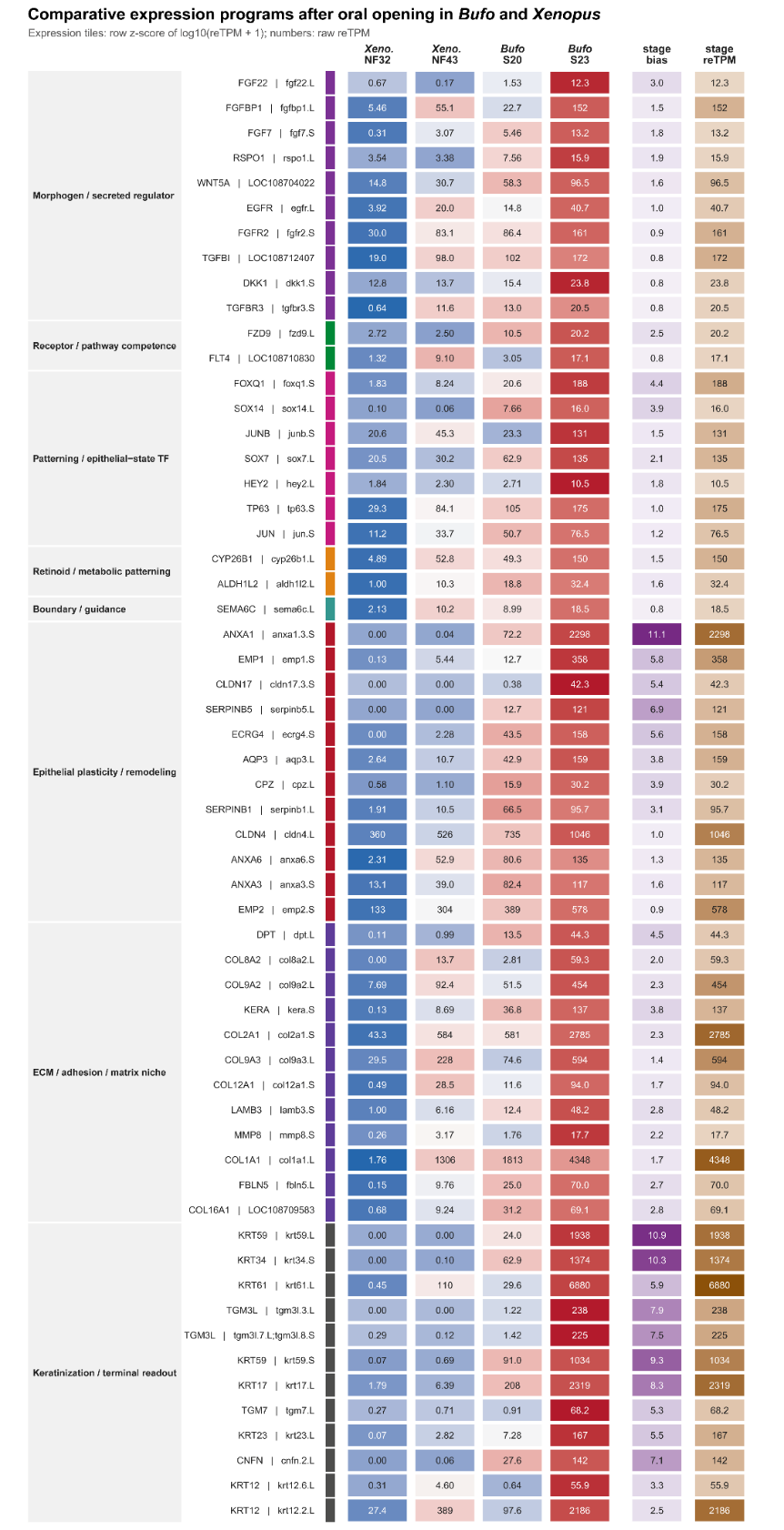

**Fig. S8 Comparative expression program after primary mouth opening in *Bufo* and *Xenopus*.**

Representative genes upregulated after oral opening are grouped by functional category and compared across Xenopus NF32/NF43 and Bufo S20/S23. Tile colors indicate row z-scores of log10 (reTPM + 1), and numbers indicate raw reTPM values. The post-opening program includes FGF, Wnt, TGFβ and EGFR pathway components, epithelial-state regulators, ECM/adhesion genes and keratinization markers, indicating activation of epithelial remodeling, matrix organization and terminal differentiation after mouth opening.

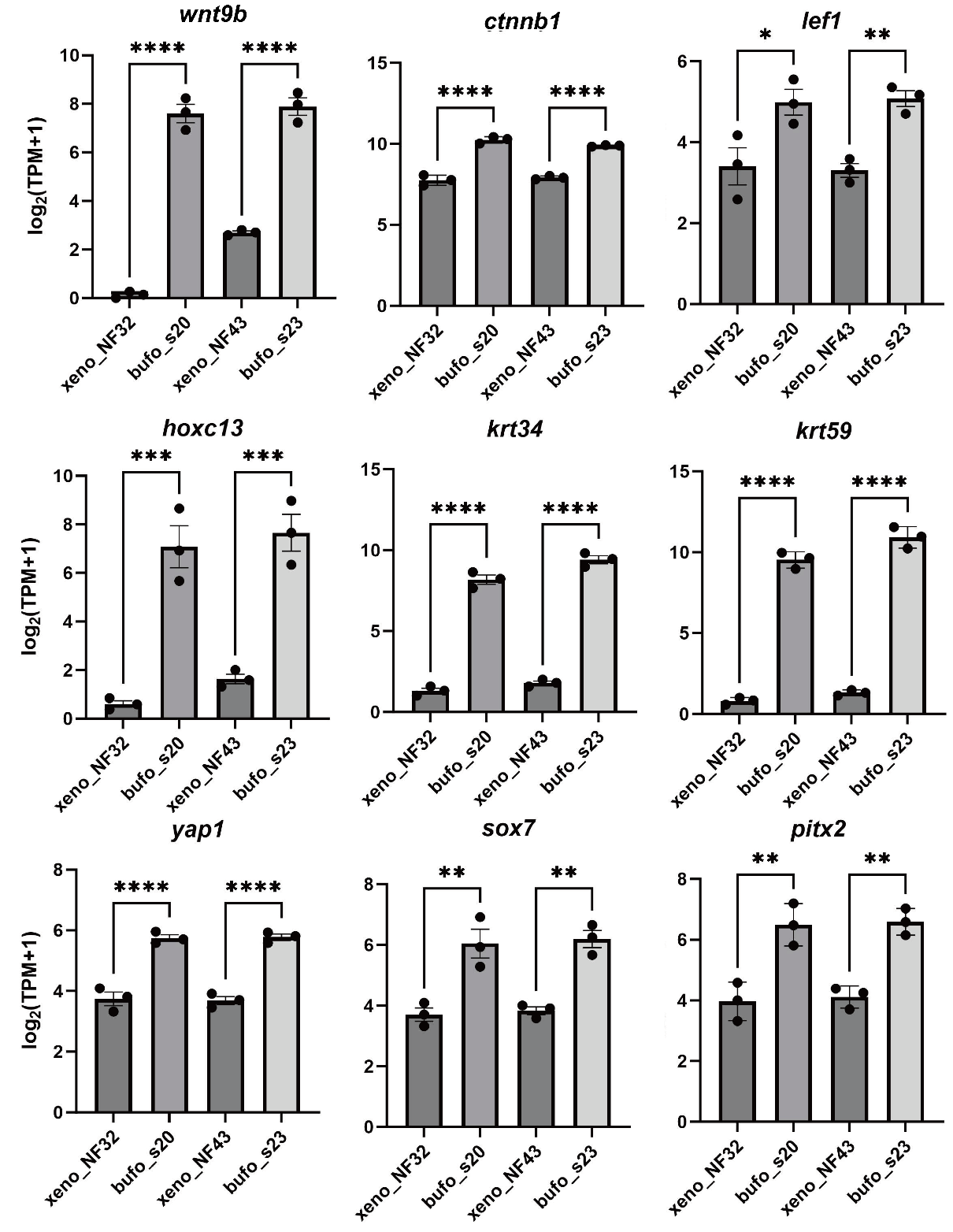

**Fig. S9 Expression comparison of selected oral-disc-associated genes between *Bufo* and *Xenopus* around primary mouth opening.**

Bar plots showing log₂ (TPM + 1) expression values of selected regulatory, signaling-associated, and keratinization-related genes in *Bufo gargarizans* and *Xenopus laevis* oral samples before and after primary mouth opening. Pre-mouth-opening stages are represented by *Bufo* S20 and *Xenopus* NF32, and post-mouth-opening stages are represented by *Bufo* S23 and *Xenopus* NF43. Bars indicate means, black dots indicate biological replicates, and error bars indicate SD. Statistical significance is indicated by asterisks above brackets. TPM, transcripts per million.

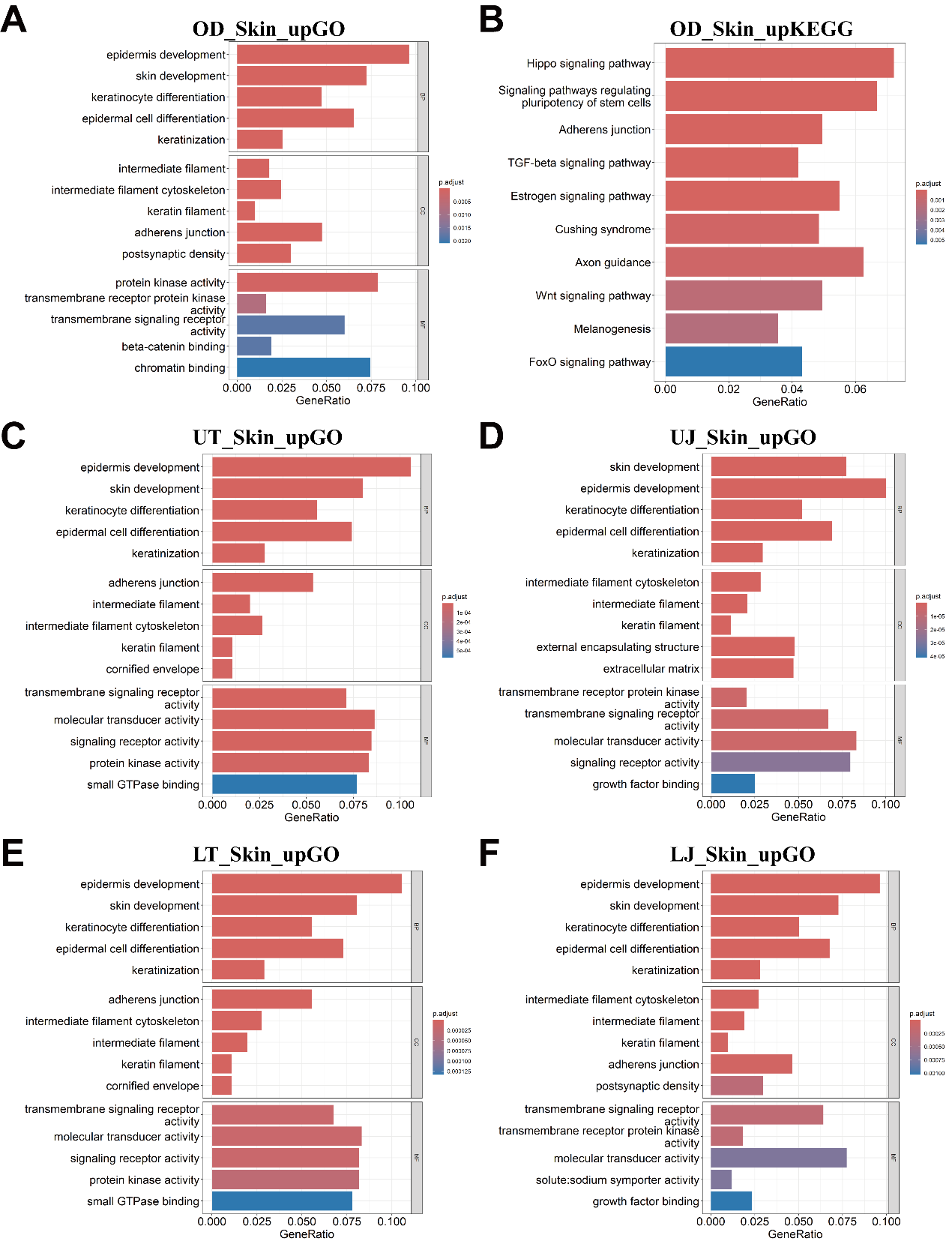

**Fig. S10 GO and KEGG enrichment analyses of genes upregulated in oral-disc tissues relative to abdominal skin at S36.**

**A**, GO enrichment analysis of genes upregulated in the whole oral disc (OD) relative to stage-matched abdominal skin at S36. **B**, KEGG enrichment analysis of genes upregulated in the whole oral disc relative to stage-matched abdominal skin at S36. **C–F**, GO enrichment analyses of genes upregulated in individual oral-disc subregions relative to stage-matched abdominal skin, including the upper labial tooth row (UT; **C**), upper jaw sheath (UJ; **D**), lower labial tooth row (LT; **E**), and lower jaw sheath (LJ; **F**). GO terms are grouped into biological process (BP), cellular component (CC), and molecular function (MF) categories. The y-axis indicates enriched GO terms or KEGG pathways, the x-axis indicates GeneRatio, and color denotes adjusted *P* values (*p. adjust*).

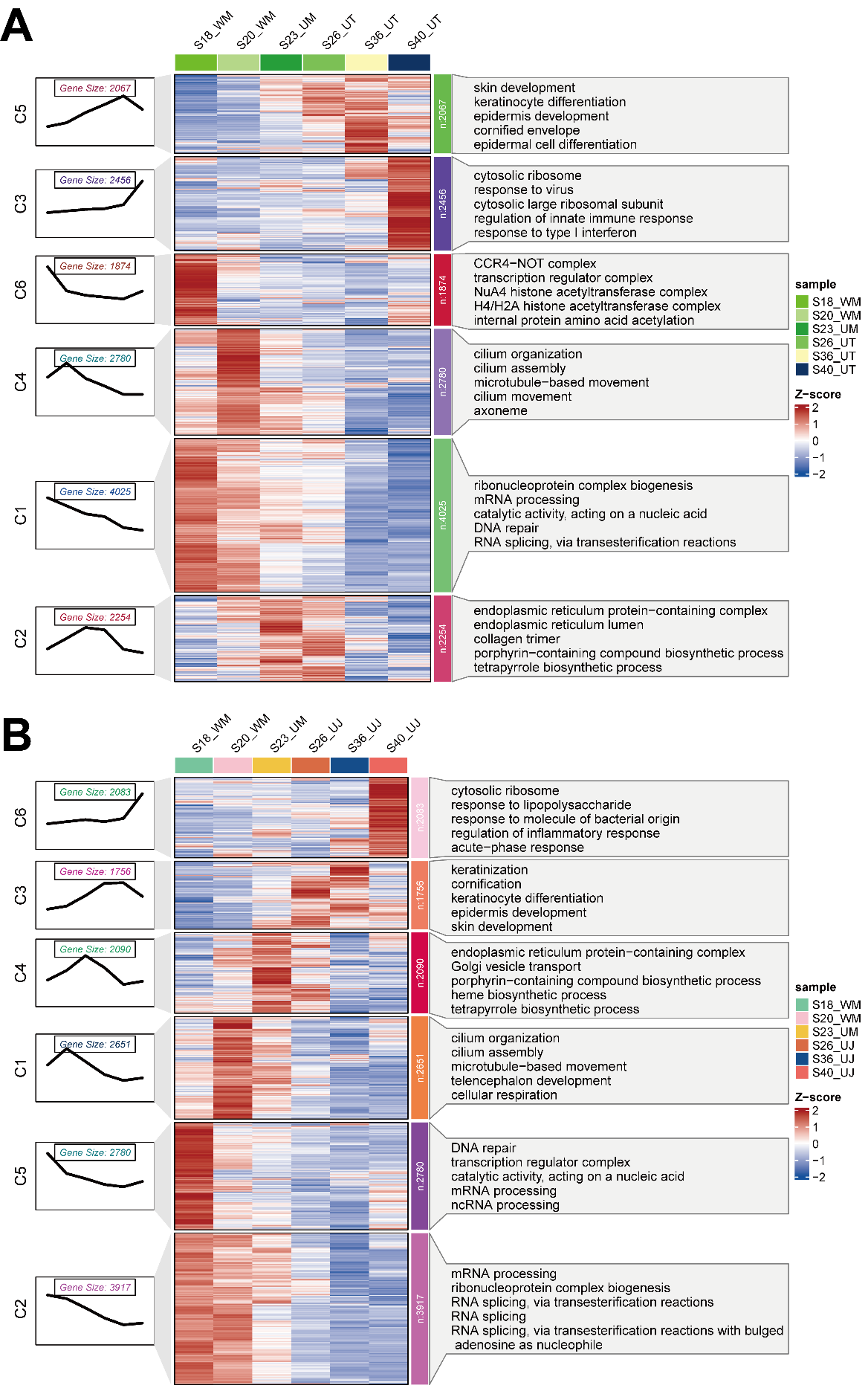

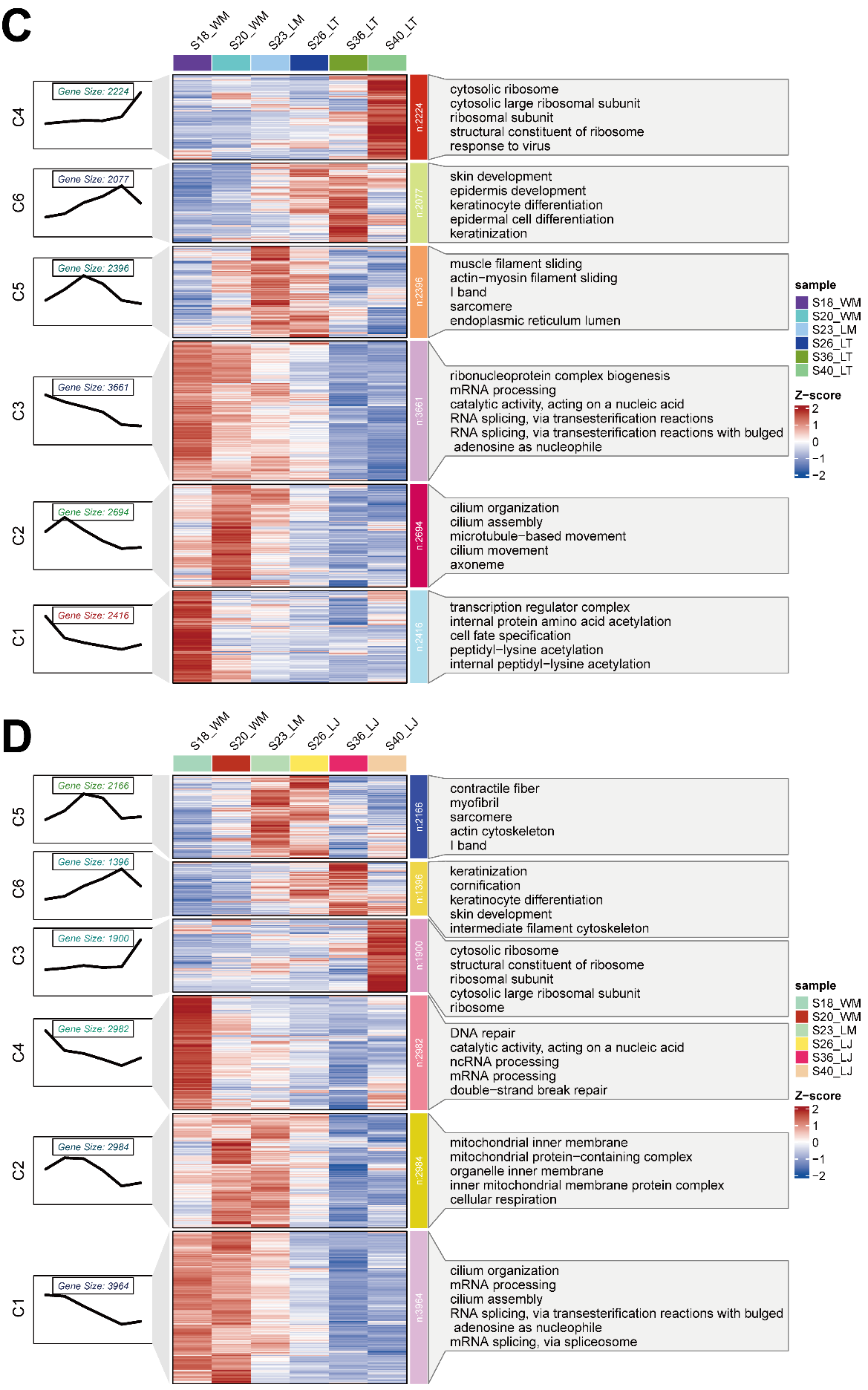

**Fig. S11** **Mfuzz clustering and GO enrichment analyses of gene-expression dynamics in oral-disc subregions of *Bufo gargarizans*.**

A-D, Mfuzz clustering of developmental transcriptomes from the upper labial tooth row (UT; A), upper jaw sheath (UJ; B), lower labial tooth row (LT; C), and lower jaw sheath (LJ; D). In each panel, line plots show the mean standardized expression profile of each gene module, heatmaps show gene-expression Z-scores across developmental samples, and the right panels show the top five enriched GO terms for each module.

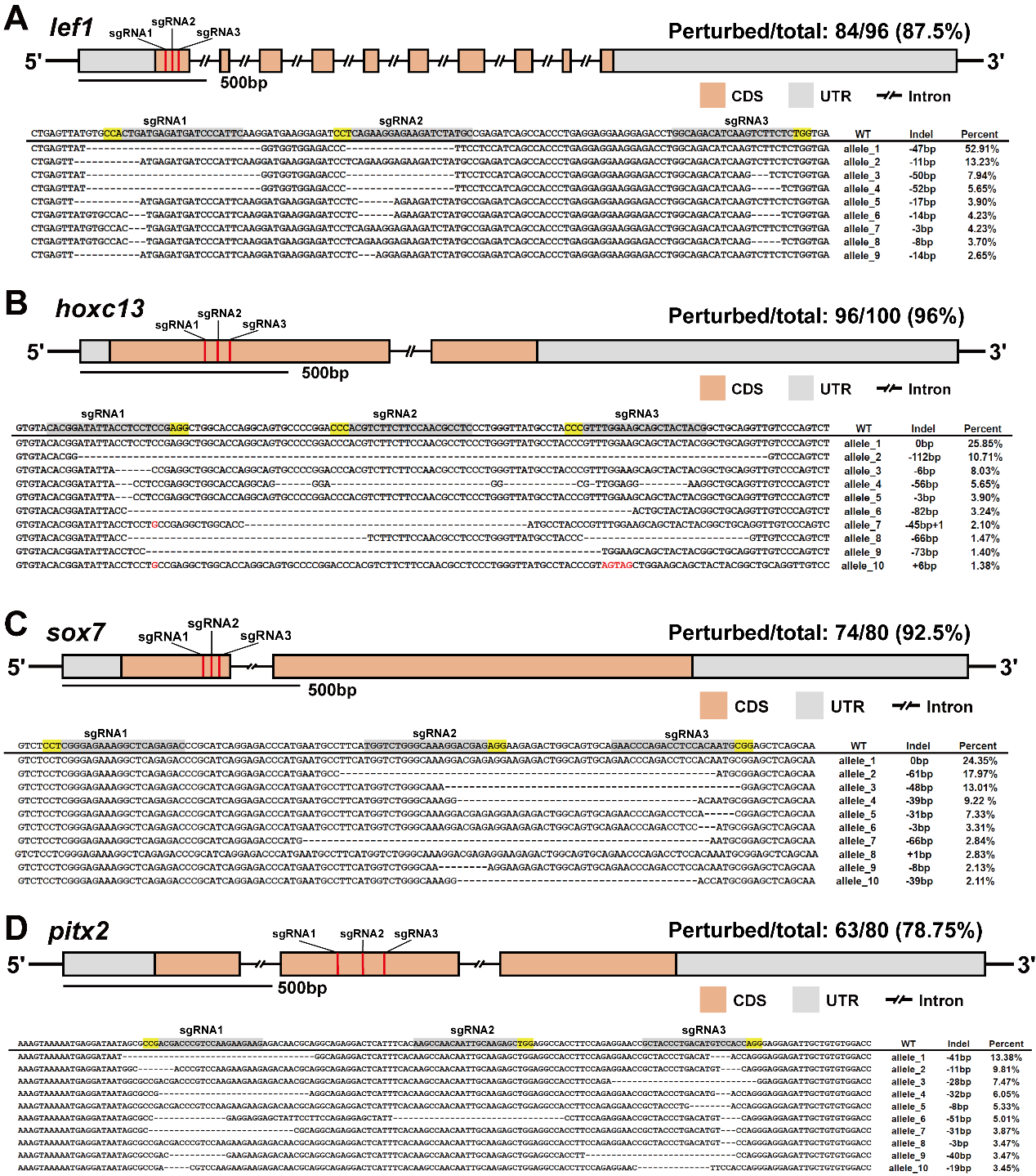

**Fig. S12** **CRISPR/Cas9 target design and Fast-NGS genotyping of representative F0 perturbed tadpoles.**

**A–D,** Target-site design and amplicon-based Fast-NGS genotyping for *lef1* (**A**), *hoxc13* (**B**), *sox7* (**C**), and *pitx2* (**D**). Gene schematics show the positions of three sgRNA target sites for each gene. Apricot boxes indicate coding sequences (CDS), grey boxes indicate untranslated regions (UTRs), and double slashes indicate omitted intronic intervals. Sequence alignments show representative amplicon reads from individual F0 edited tadpoles, with the wild-type reference sequence shown at the top. Grey shading marks sgRNA target regions, yellow highlighting marks PAM sequences, dashes indicate deleted nucleotides, and red letters indicate inserted or substituted nucleotides.

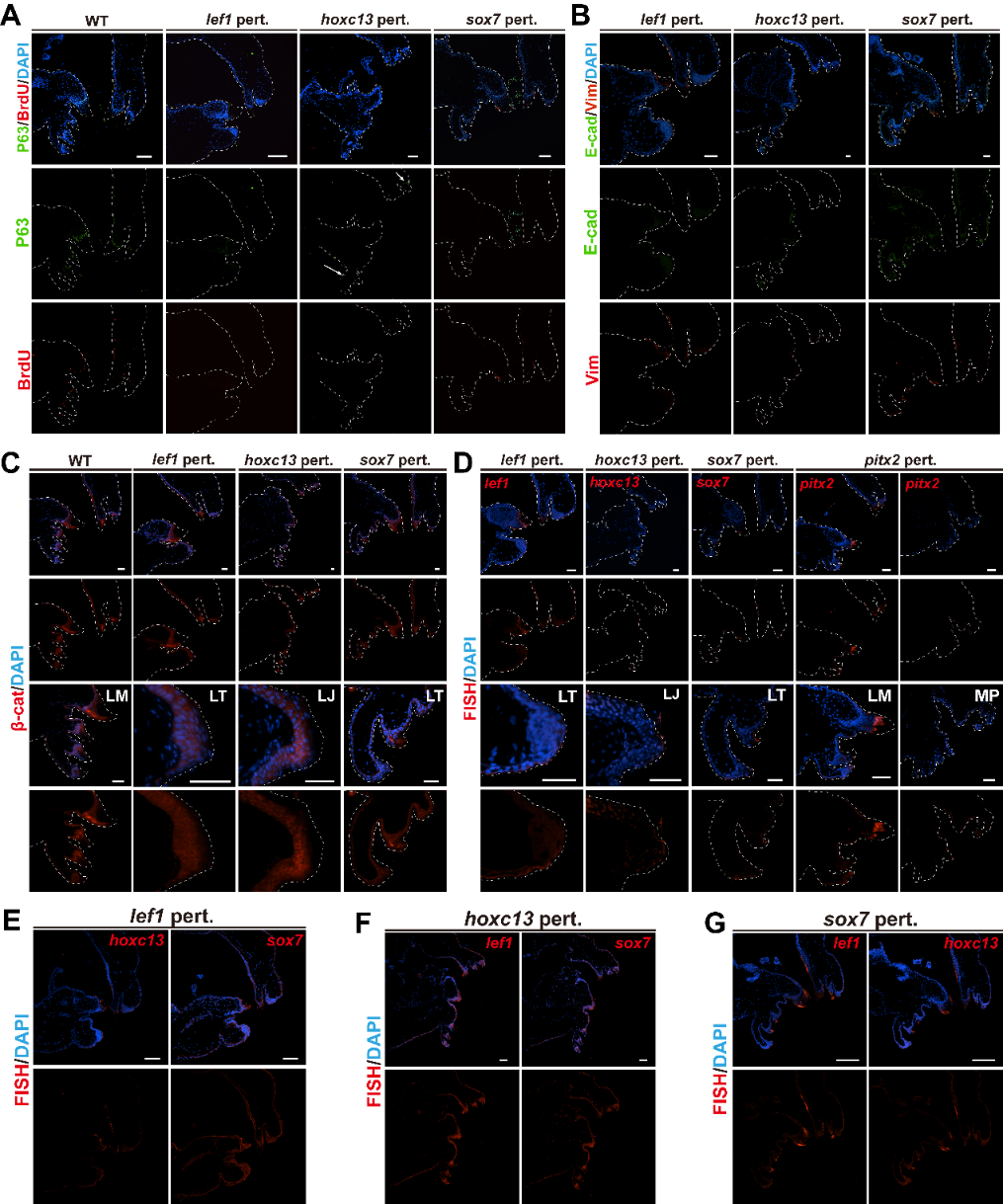

**Fig. S13 Full-field images of immunofluorescence and HCR RNA-FISH after selected-gene functional perturbation.**

Full-field images corresponding to the representative regions shown in Figure 4. Dashed lines delineate oral-disc structures. **A,** p63 and BrdU immunofluorescence in wild-type (WT) and *lef1*-, *hoxc13*-, and *sox7*-perturbed oral discs. Upper panels show merged p63, BrdU, and DAPI signals; lower panels show p63 and BrdU signals separately. **B,** E-cadherin and vimentin immunofluorescence in *lef1*-, *hoxc13*-, and *sox7*-perturbed oral discs. Upper panels show merged E-cadherin, vimentin, and DAPI signals; lower panels show E-cadherin and vimentin signals separately. **C,** β-catenin immunofluorescence in WT and *lef1*-, *hoxc13*-, and *sox7*-perturbed oral discs. Full-field views and enlarged regions are shown for each condition. **D,** HCR RNA-FISH analysis of target-gene expression in *lef1*-, *hoxc13*-, *sox7*-, and *pitx2*-perturbed oral discs. Full-field views and enlarged regions show the corresponding HCR signal and DAPI counterstaining. **E,** HCR RNA-FISH analysis of *hoxc13* and *sox7* expression in *lef1*-perturbed oral discs. **F,** HCR RNA-FISH analysis of *lef1* and *sox7* expression in *hoxc13*-perturbed oral discs. **G,** HCR RNA-FISH analysis of *lef1* and *hoxc13* expression in *sox7*-perturbed oral discs. **Scale bars = 50 μm in A-D. Scale bars = 100 μm in E-G.**

**Abbreviations:** E-cad, E-cadherin; LJ, lower jaw region; LM, lower mouth region; LT, lower tooth-row region; MP, marginal papillae; Vim, vimentin; WT, wild type.

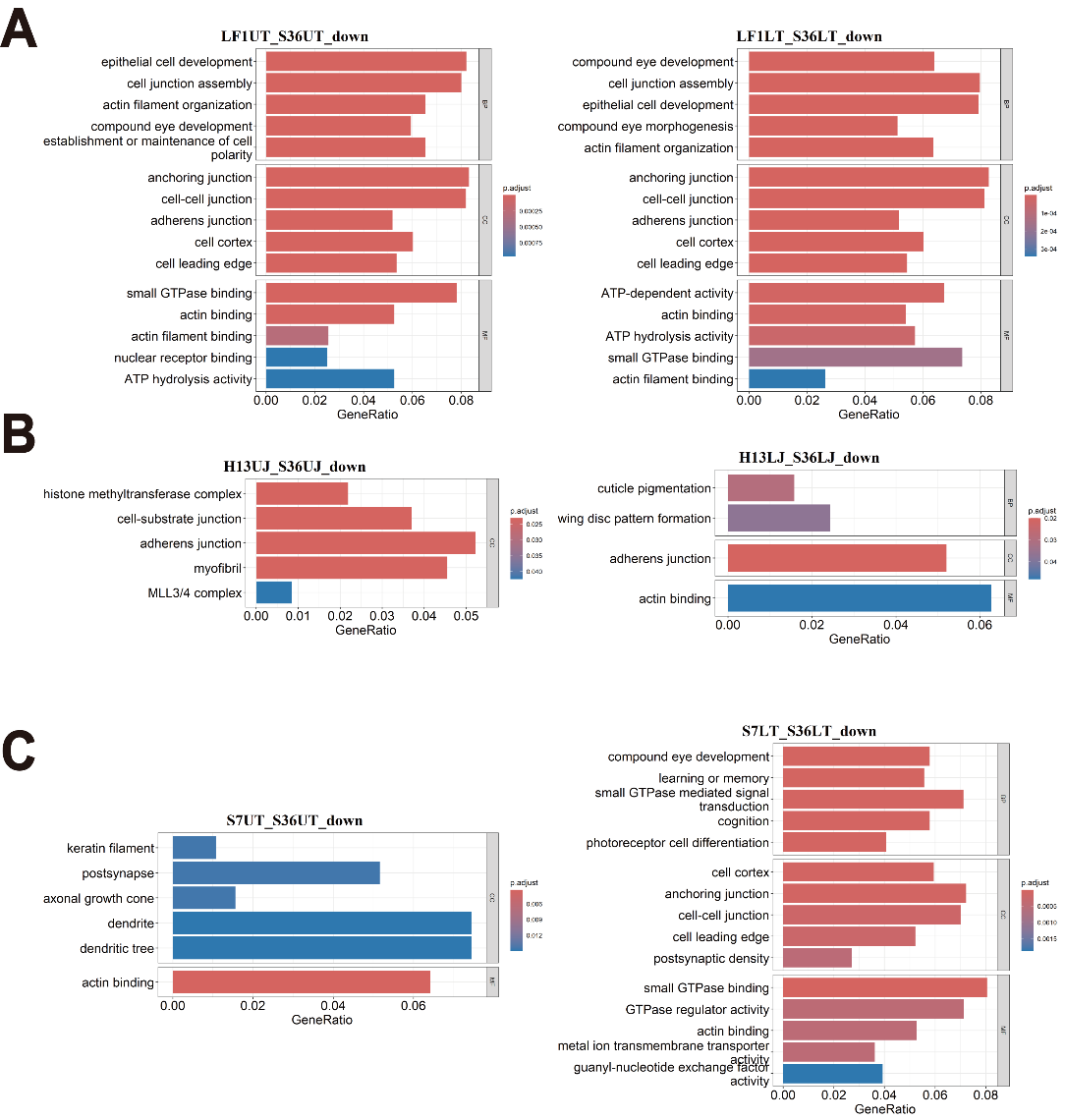

**Fig. S14 GO enrichment analysis of genes downregulated after selected-gene functional perturbation in S36 oral-disc subregions.**

**A–C,** GO enrichment analysis of genes downregulated in selected oral-disc subregions after functional perturbation relative to the corresponding wild-type S36 subregions. **A,** Enriched GO terms for genes downregulated in the upper tooth-row (UT) and lower tooth-row (LT) regions after *lef1* functional perturbation. **B,** Enriched GO terms for genes downregulated in the upper jaw (UJ) and lower jaw (LJ) regions after *hoxc13* functional perturbation. **C,** Enriched GO terms for genes downregulated in the UT and LT regions after *sox7* functional perturbation. Facets indicate biological process (BP), cellular component (CC), and molecular function (MF) categories; categories without significant enrichment are not shown. Bar length indicates GeneRatio, and bar color indicates adjusted *P* value.

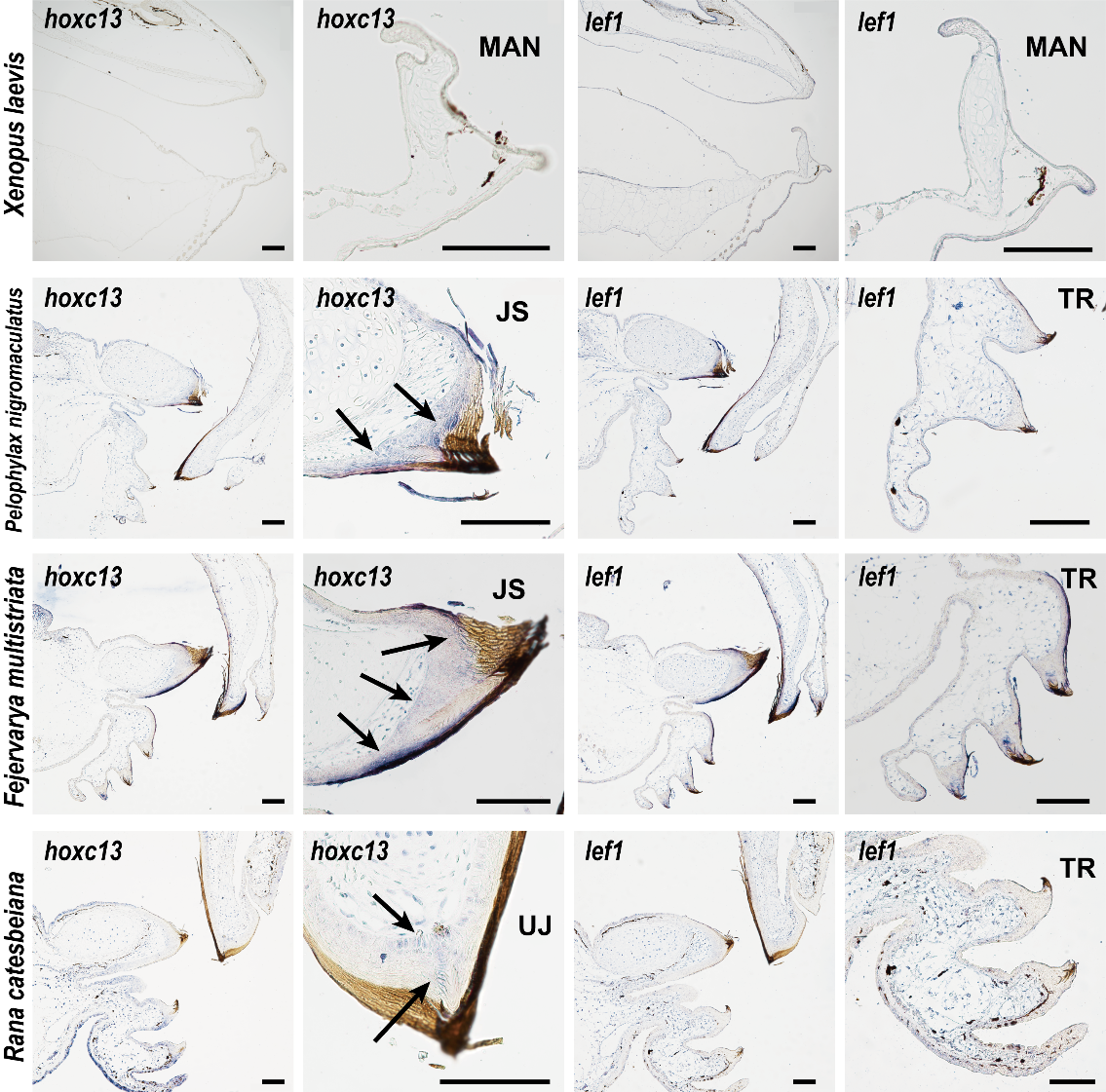

**Fig. S15 *in situ* hybridization of *hoxc13* and *lef1* in tadpole oral structures across selected anuran species.**

Representative RNA in situ hybridization images showing *hoxc13* and *lef1* expression in larval oral regions of *Xenopus laevis*, *Pelophylax nigromaculatus*, *Fejervarya multistriata*, and *Rana catesbeiana*. In *X. laevis*, which lacks a typical keratinized oral disc, comparable regionalized *hoxc13* or *lef1* expression domains were not observed. In species with typical oral discs, *hoxc13* signal was enriched in jaw-sheath regions, whereas *lef1* signal was associated with tooth-row regions. Black arrows indicate *hoxc13*-positive domains in jaw-sheath regions. **Scale bars = 150 μm.**

**Abbreviations:** JS, jaw sheath; MAN, mandibular cartilage; TR, tooth row; UJ, upper jaw sheath.

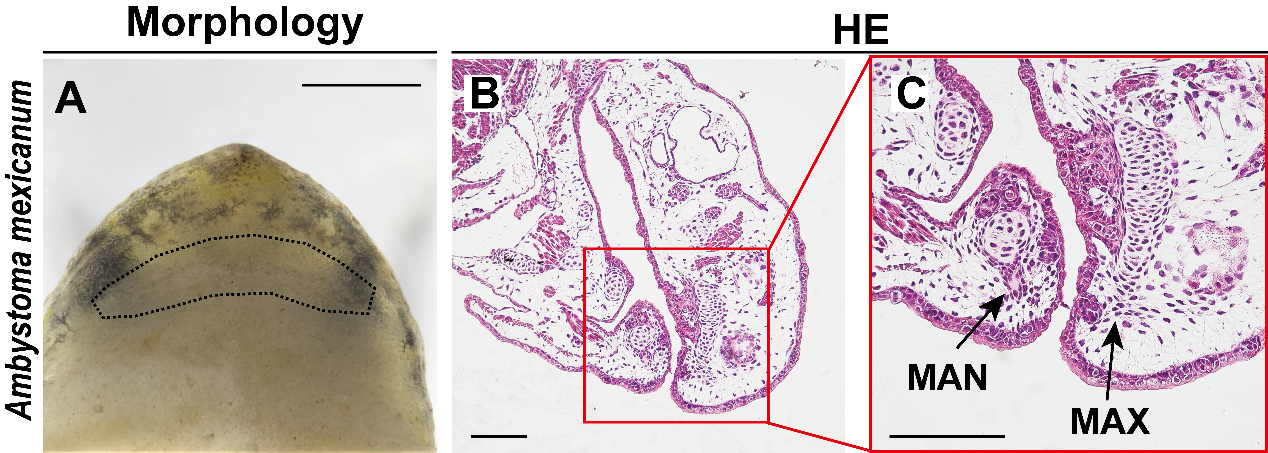

**Fig. S16 Larval oral morphology and histology of *Ambystoma mexicanum*.**

**A,** Gross morphology of the larval mouth region in *Ambystoma mexicanum*. Dashed lines indicate the mouth opening. **B–C,** H&E-stained sections of the larval oral region, with the boxed area in **B** enlarged in **C**. The larval mouth lacks typical anuran oral-disc-associated structures, including keratinized jaw sheaths and labial tooth rows, and is supported by maxillary and mandibular cartilages. **Scale bars: morphology = 1,000 μm; H&E = 250 μm.**

**Abbreviations:** MAN, mandibular cartilage; MAX, maxillary cartilage.

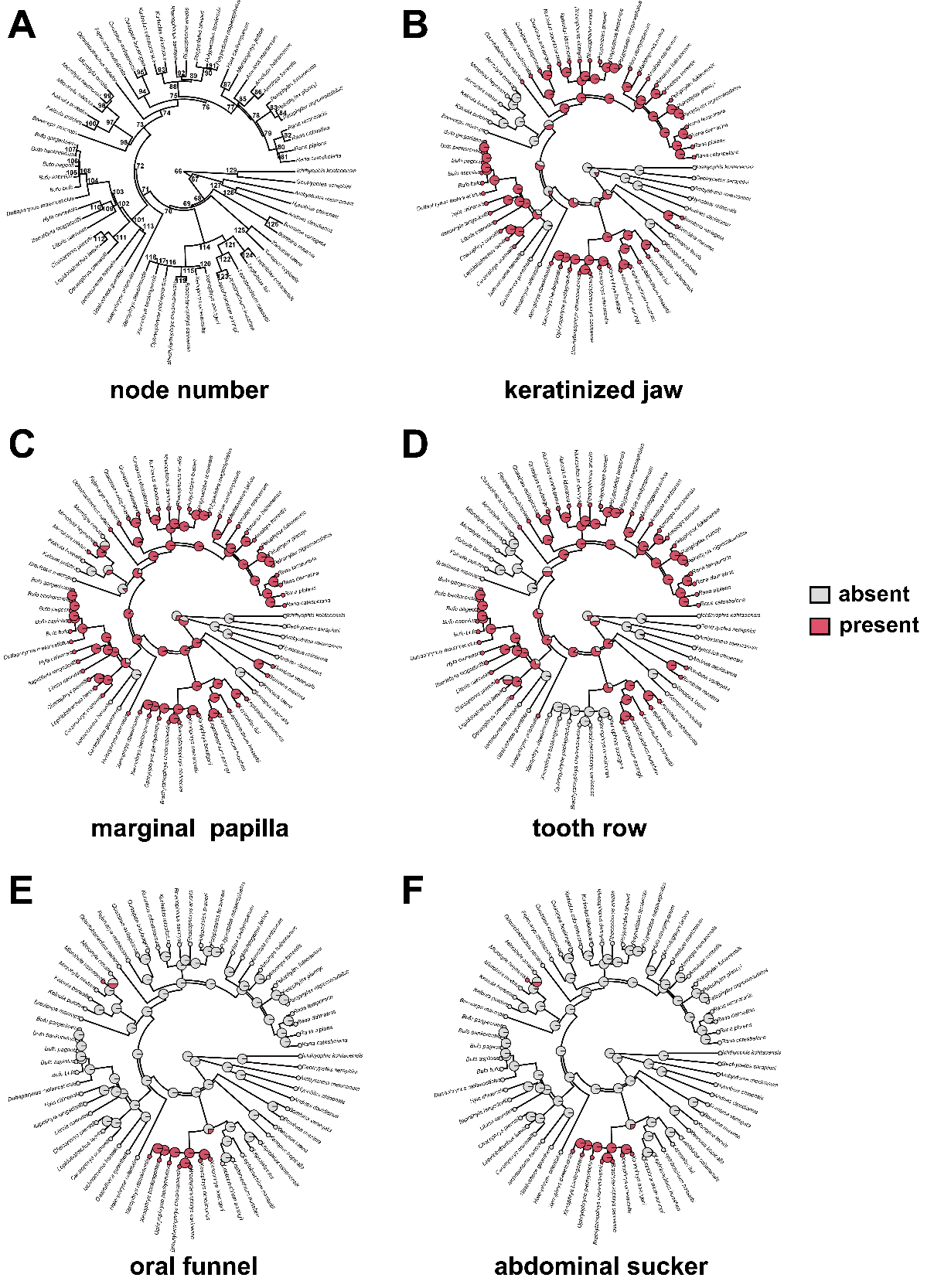

**Fig. S17 Node-labelled phylogeny and ancestral-state reconstructions of larval oral structural characters.**

**Node-labelled phylogeny and ancestral-state reconstructions of larval oral structural characters.**
**A,** Phylogenetic tree with internal-node IDs used for cross-reference in the text. **B–F,** Binary ancestral-state reconstructions for jaw sheath (**B**), marginal papillae (**C**), tooth rows (**D**), oral funnel (**E**), and abdominal sucker (**F**). Tip states were scored from direct observations and literature records, as summarized in Table S2. Internal nodes are colored according to the marginal posterior probability of character presence on a 0–1 scale. Reconstructions support ancestral presence of jaw sheaths and tooth rows, repeated gains of marginal papillae, restricted occurrence of oral funnels, and limited retention of abdominal suckers across sampled lineages. Reconstructions were performed under an Mk model with an ER transition structure. Taxa lacking a tadpole stage were coded as absence for larval oral structural characters.
